# A method for chronic and semi-chronic microelectrode array implantation in deep brain structures using image guided neuronavigation

**DOI:** 10.1101/2022.08.26.505452

**Authors:** Borna Mahmoudian, Hitarth Dalal, Jonathan Lau, Benjamin Corrigan, Kevin Barker, Adam Rankin, Elvis C.S. Chen, Terry Peters, Julio C. Martinez-Trujillo

## Abstract

Precise targeting of deep brain structures in humans and large animal models has been a challenge for neuroscientists. Conventional protocols used in animal models typically require large access chambers which are prone to infection and involve assembly and implantation of complex microdrives for semi-chronic applications. Here we present a methodology for improving targeting of subcortical structures in large animals such as macaque monkeys, using image guided neuronavigation. Design of custom cranial caps allowed for incorporation of stable fiducial markers, required for increased targeting accuracy in neuronavigation procedures, resulting in an average targeting error of 1.6 mm over three implantations. Incorporation of anchor bolt chambers, commonly used in human neurosurgery, provided a minimally invasive entrance to the brain parenchyma, allowing for chronic recordings. By leveraging existing 3D printing technology, we fabricated an anchor bolt-mounted microdrive for semi-chronic applications. Our protocol leverages commercially available tools for implantation, decreases the risk of infection and complications of open craniotomies, and improves the accuracy and precision of chronic electrode implantations targeting deep brain structures in large animal models.

## Introduction

Experimental animal models are an indispensable tool in biomedical research aiming to uncover the neural basis of behavior and disease. Non-human primates’ (NHP) strong brain homology with humans and similarities in behavioural repertoire make them a suitable model for studying neural circuits involved in cognitive and emotional processes (Bernardi & Salzman, 2019). Furthermore, NHPs have served as models for developing techniques and methodologies that can improve procedures used in medical practice. One such example has been their critical contribution to the development of deep brain electrical stimulation techniques currently used in the treatment of neurological diseases such as Parkinson Disease (Benazzouz et al., 1993). Consequently, in vivo extracellular recording of neural activity in behaving NHPs can provide important insights into the neural correlates of human behaviour and disorder. Factors in consideration for an in vivo extracellular implantation include methodologies for targeting the regions of interest and the recording technique (i.e., chronic, semi-chronic, or acute). The methodology for targeting the regions of interest influences the implantation accuracy (Walbridge et al., 2006), and the recording technique determines the duration of the electrode implant in the brain i.e., for hours, a single day or across days (McMahon et al., 2014). While protocols for targeting and recording from the cortical surface have been described in previous literature (Sponheim et al., 2021), precise subcortical targeting and recording techniques are technically more challenging and therefore reported more scarcely.

Targeting of the cortical surface benefits from visual access to the sulci and gyri that delineate the areas of interest, which along with a generalized stereotactic atlas or the subject’s MRI (Premereur et al., 2020) can provide accurate targeting. In contrast, implantation of deep-brain structures remains a challenge, as small angular deviations in the trajectory of the implantation can lead to significant targeting errors given the depth and the geometry of the implantations (Bjartmarz & Rehncrona, 2007). Protocols for chronic, semi-chronic, and acute recordings have been well established for cortical surface implantations in both humans and NHPs (Ferrea et al., 2018; Mountcastle et al., 1991; Paulk et al., 2022; Sponheim et al., 2021). Although similar recording protocols for implantation of subcortical structures exist in humans (Patel et al., 2013), chronic and semi-chronic approaches for subcortical recordings in large experimental animals such as NHPs are limited. Existing protocols commonly involve fabrication and implantation of custom-made recording chambers and assembly of complex microdrive systems (McMahon et al., 2014; Paulk et al., 2022). Furthermore, such recording chambers usually require an open craniotomy that requires daily cleaning under sterile conditions, and opens the door for opportunistic infections that can jeopardize the health of the animal, implant, and the results of the experimental manipulations and procedures.

In this paper we aim to address the aforementioned challenges by first developing a targeting methodology for implantation of electrodes in subcortical structures, and second by developing a chronic and semi-chronic technique for recording from the implanted deep-brain regions in NHPs. To this end, we have utilized a neuronavigation toolkit (Brainsight, Rogue Research Inc) to establish an accurate and precise, subject-specific, deep-brain implantation targeting protocol. Additionally, adapting anchor bolt (Ad-Tech Medical) chambers from human neurosurgery has allowed for a minimally invasive electrode array implantation of amygdalae of two NHP subjects. Computed tomography (CT) in vivo imaging established an average Euclidean targeting error of 1.6 mm over three implantations on two subjects (subject Mi: 1.2 mm and 1.4 mm, subject Ma: 2.3 mm), and viable recordings were obtained for a period of at least four months. We will first provide a brief overview of existing methodologies in the field, highlighting the advantages of our approach. Our targeting methodology and the implantation protocol will then be described, with the resulting targeting error and recording samples presented thereafter. We finally will provide suggestions for future improvement of targeting methodologies.

### Overview of Subcortical Targeting Methodologies

The traditional gold standard method for electrode implantation in deep-brain structures in NHPs has been the frame-based stereotaxic method (Saunders et al., 1990). In this procedure, implantation trajectories are determined by locating anatomical landmarks (e.g., bony landmarks such as the ear canals). Referencing these landmarks on a generalized stereotactic atlas, or in pre-operative images of the animal aligned within a stereotaxic frame, allows for establishing of the implantation trajectories. Despite the extensive use of this method, there are several disadvantages: 1) If pre-operative imaging is used, misalignment of the animal within the stereotaxic frame during the operation compared to the pre-operative images can lead to targeting errors (Walbridge et al., 2006), 2) the frame is bulky, restricting access to certain regions (Frey et al., 2004), and the range of motion available when using a frame prevents certain procedures (Grimm et al., 2015), 3) The use of a standard stereotactic atlas for determination of implantation coordinates and trajectories can lead to targeting errors, as these atlases are derived from averages of several brains, ignoring individual subject variability (Bickart et al., 2011; Sallet et al., 2011). 4) As the trajectories are determined pre-operatively, this leads to inflexibility of trajectory adjustment during surgery.

To address these concerns, we developed a surgical protocol using a frameless stereotaxic system (also referred to as neuronavigation), which addresses many of the problems with frame-based approaches. The accuracy and precision of the latest frameless systems in human neurosurgery have reached and likely surpassed that of frame-based methods (Dhawan et al., 2019; Holloway et al., 2005; Li et al., 2016; Roth et al., 2018). This approach allows surgical flexibility by reducing bulk of the surgical frame, and the use of computer-assisted technologies which provide real-time feedback (Frey et al., 2004). Importantly, target coordinates and trajectories are determined using the subject’s MRI as opposed to a generalized atlas (Frey et al., 2004), allowing for both a smaller craniotomy for gyri/sulci assessment, as well as accounting for individual differences in subcortical anatomy (Bickart et al., 2011; Sallet et al., 2011; Saunders et al., 1990). To avoid invasive procedures, previous protocols (Johnston et al., 2016) have used dental imprint platforms or skin markers as fiducials (analogous to boney landmarks used in stereotaxic surgeries) for registration of the subject-scan space. The use of such fiducials typically leads to a low registration accuracy as they are difficult to repeatedly localize, prone to movement between image acquisition and surgery, and are improperly configured or distributed around the target of interest (Holloway et al., 2005; Wang & Song, 2011). We sought to improve the previous protocols by fabricating a custom cranial cap for each subject (Blonde et al., 2018), and utilizing the cranial cap screws as registration fiducials for the neuronavigation procedure. This allowed us to incorporate fiducial markers, a headpost mount, and housing for electrophysiological equipment into a single biocompatible design.

### Overview of Subcortical Recording Techniques

Electrophysiological recording techniques can be categorized based on the duration of the electrode implant in the brain. Acute implants consist of recording electrodes inserted only for the duration of a daily recording session and subsequently removed. While this technique is effective and flexible in spatially sampling an anatomical region of interest (Morrow et al., 2020), it requires daily time-consuming steps such as insertion of electrodes, pre-and-post implant cleaning, as well as having an increased susceptibility to infection (Dotson et al., 2017). Moreover, they do not allow for longitudinal recordings of the same population of neurons across sessions. In contrast, a chronic implant is positioned permanently inside the brain for the during of the experimental period (e.g. months), allowing for longitudinal monitoring of neurons while reducing preparation time and limiting tissue exposure to potential pathogens (McMahon et al., 2014). However, as postoperative repositioning of the electrode is not possible, the tissue’s foreign body response could develop, reducing the viability of the implant as the signal quality degrades (Groothuis et al., 2014; Kozai et al., 2015). A semi-chronic implant is a hybrid approach that addresses some of the limitations of both acute and chronic methods (Dotson et al., 2017). This technique requires a microdrive, which allows for movement of an electrode along a single axis. This enables both longitudinal recording of a neuronal population across sessions, as well as the ability to reposition the electrode post-operatively, yielding a new set of neurons and delaying the adverse effects of the foreign body response on the implantation (Dotson et al., 2017; Kozai et al., 2015; McMahon et al., 2014). The current availability of precise and affordable 3D printing technology allows for the design and development of custom microdrives (Headley et al., 2015). However, protocols for chronic and semi-chronic subcortical recording techniques are scarce, with the existing methods requiring fabrication and implantation of custom-made chambers, and assembly of complex microdrives. Moreover, current methods typically require large craniotomies and bulky chambers that do not allow implantation of several adjacent trajectories with different angles of approach. Here we have utilized Ad-Tech Medical anchor bolts as an electrode chamber for chronic recordings, with an in-house fabricated manual microdrive mounted for semi-chronic recordings. The advantage of the anchor bolt is the small craniotomy needed for insertion and electrode implantation, in addition to a reduced footprint which allows for implantation of multiple adjacent trajectories in nearby regions with differing approach angles. As the anchor bolt is inserted through the skull and positioned above the dura, it allows for hermetic isolation of the craniotomy from the extracranial space, reducing the chance of infection from the outside environment. Lastly, this implantation toolkit is commercially readily available from Ad-Tech Medical, making the method accessible to investigators across many laboratories.

## Materials & Methods

### Ethics Statement

As previously detailed elsewhere (Roussy et al., 2021) animal care and handling including basic care, animal training, and surgical procedures were pre-approved by the University of Western Ontario Animal Care Committee. This approval ensures that federal (Canadian Council on Animal Care), provincial (Ontario Animals in Research Act), regulatory bodies (e.g.: CIHR/NSERC), and other national Canadian Association for Laboratory Animal Medicine (CALAM) standards for the ethical use of animals are followed. Regular assessments for physical and psychological well-being of the animals were conducted by researchers, registered veterinary technicians, and veterinarians.

### General workflow

The general workflow of the procedures is presented in Figure 1. In the different sections we will refer to the components of this flowchart. Inventory of all the tools and software required for replication of this method are presented in Supplementary Table 1.

**Figure 1:**
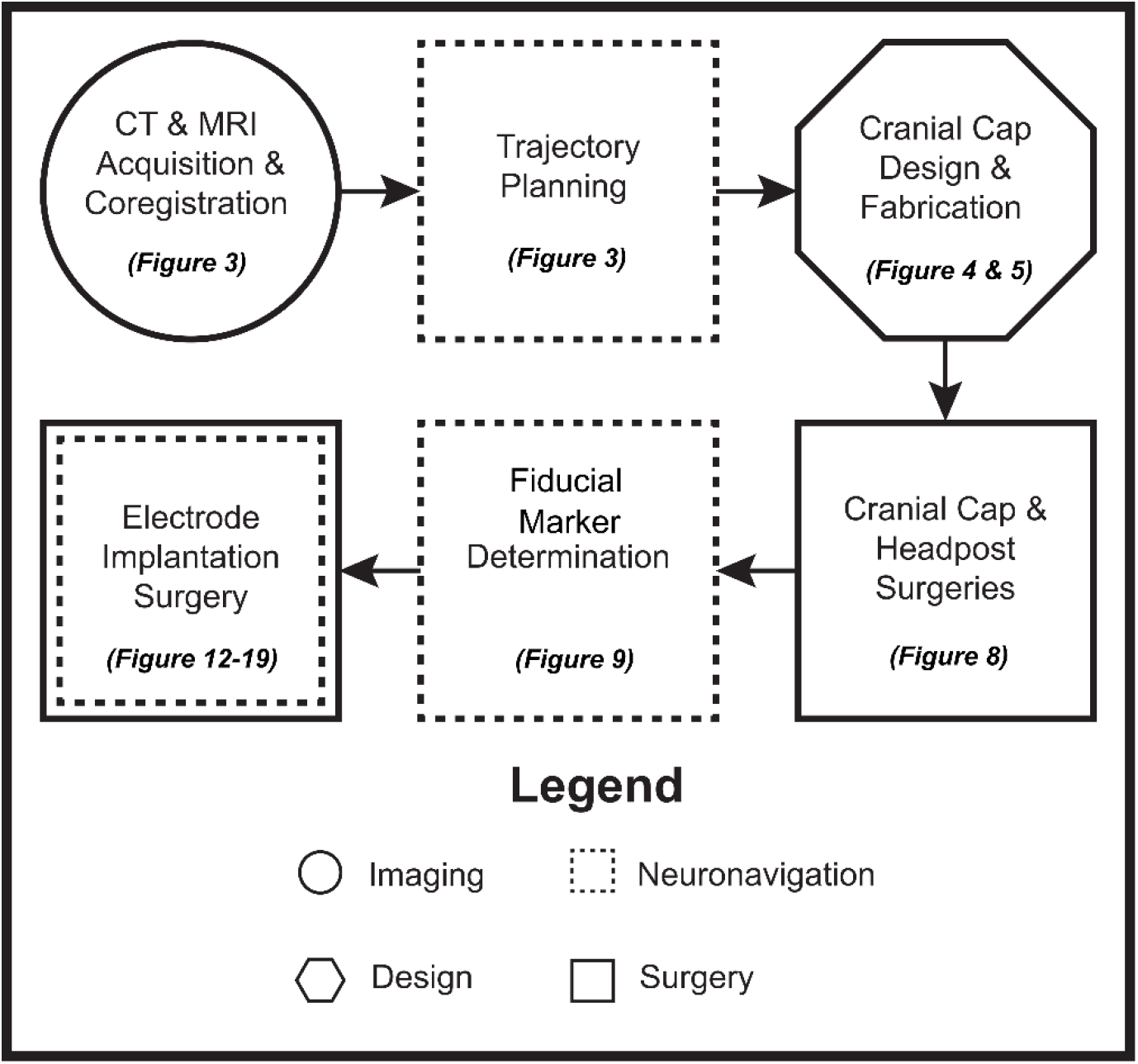
Flowchart presenting an overview of the electrode implantation method.

### Electrode apparatus assembly

#### Microwire Brush Array (MBA)

The MBA (Microprobes for Life Science, Gaithersburg, MD) consists of a bundle of 64 or 32 microwires (12.5 µm diameter each) encased by a microfil tube and an external ground wire (Fig. 2A). The microwires extend beyond the microfil and splay to allow for the recording of a population of neurons. Microwire brush arrays have been previously used to target deep-brain structures in NHPs (Bondar et al., 2009; McMahon et al., 2014) yielding longitudinal recordings of single units. While the implanted electrode in this procedure is the MBA, we believe that the described method will be extendible to other single shank recording electrodes, taking into consideration the diameter of the access chamber.

**Figure 2:**
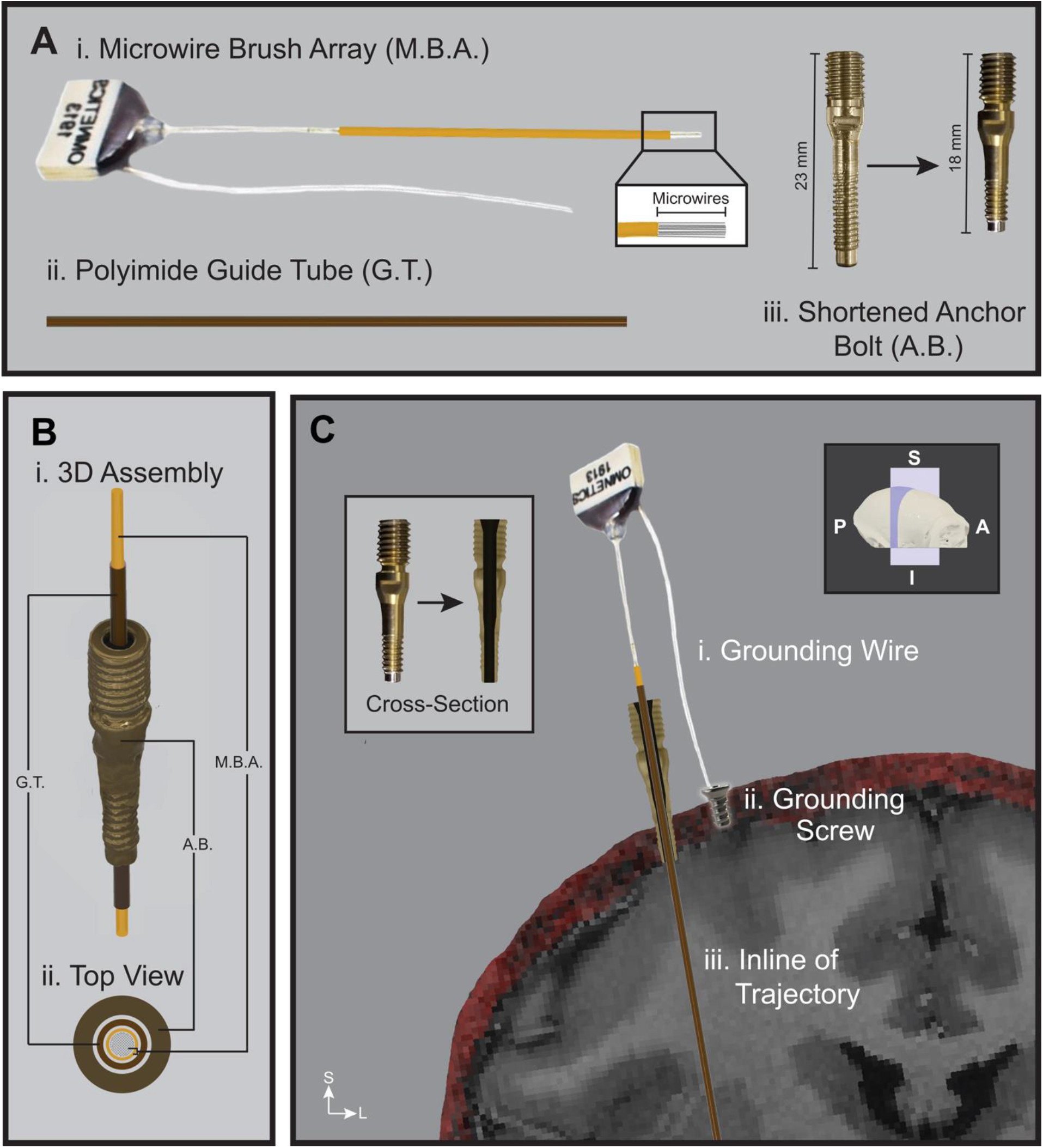
Hardware tools and assembly for the electrode implantation. **A.** Components of the electrode implantation assembly*. (i) Microwire Brush Arrays (MBA): 64 or 32 microwires extend 2 or 5 mm beyond the microfil tube. (ii) Polyimide guide tube. (iii) Anchor Bolt: A chamber for the electrode. Shortened to accommodate the thinner NHP skull. **B.** 3D reconstruction of the assembly components*: (i) Angled view shows the insertion of electrode and guide tube into the anchor bolt. (ii) Top view is a cross-section of the assembly at the top of the anchor bolt. **C.** Schematic of assembled implantation components on a coronal section of the subject’s MRI and CT (red)*: (i) The grounding wire will be attached to the (ii) grounding screw. (iii) The entire assembly will be fixed in-line of the desired trajectory. *Left Inset:* A coronal cross-section of the anchor bolt. *Right Inset:* The illustrated coronal plane (purple) on the subject’s skull. *Note: Dimensions are not to scale. For demonstration purposes only. Microbrush array electrode image used is modified from Microprobes For Life Science with permission. S = Superior; I = Inferior; A = Anterior; P = Posterior; L = Left; R = Right.

#### Anchor Bolt

The anchor bolt (LSBK2-BX-04, Ad-Tech Medical Instruments, Oak Creek, WI) is a minimally invasive recording chamber (2.4 mm diameter craniotomy) that is commonly used in human neurosurgery for depth electrode placement in stereoencephalography (sEEG) procedures. The chosen anchor bolts’ inner diameter was 0.99 mm while the outer diameter measured 3.96 mm, providing a low footprint on the skull. It consists of two threaded regions, a nub on the distal end, a lumen and depressions on the sides of the proximal end (Fig. 2A). The anchor bolt includes a cap that is fastened to secure electrodes used in sEEG procedures and has been removed for the current protocol. A proprietary placement wrench is inserted inside the lumen to fasten the distal threads of the anchor bolt into the skull. The nub on the distal end prevents the threads from coming in contact with the dura. Lastly, the depressions along the side provide a surface for a wrench tool when removing the bolt. Given the smaller size of a NHP subject and a thinner skull compared to humans, the distal end of the anchor bolt was reduced in length and resulted in a shorter assembly that is easier to house (Fig. 2A).

#### Guide Tube

The polyimide guide tube (MicroLumen Inc, Oldsmar, FL) provided a safe way of loading the MBA into the anchor bolt, preventing damage to the microwires (Fig. 2A). Additionally, it provided a trajectory for the electrode as it traveled towards the target, preventing deviations from the path.

#### Assembly

During the assembly, the anchor bolt was first fastened into the skull using the placement wrench. The guide tube was passed through the bolt and secured to the anchor bolt using dental cement (BISCO Dental Products, Schaumburg, Il). Finally, the MBA was passed through the guide tube and lowered to the target (Fig. 2B). The MBA was secured to the anchor bolt using dental cement and the grounding wires were wrapped around screws fastened into the skull (Fig. 2C).

### Trajectory Planning & Validation

Surgical planning began by importing imaging data into the Brainsight software (Rogue Research Inc., Montréal). Computer Tomography (CT) and Magnetic Resonance Imaging (MRI) images were acquired of the NHP subjects. Two male macaque (subject Mi, 9 year old rhesus macaque and subject Ma, 7 year old cynomolgus macaque) were used for this study. For the purposes of this study, which presents a novel electrode implantation methodology and recording technique, sex is not a variable of relevance. MRI scans included a T1 and Time of Flight (TOF) sequences (Fig. 3A). The CT was used to determine skull thickness and contour while the T1 MRI sequence was used to identify brain structures. The TOF MRI sequence allowed for the identification of major cerebral vasculature to be avoided. The CT and MRI anatomical scans were co-registered to the same image space using a rigid-body registration within the General Registration (BRAINS) module in 3D Slicer software (Fedorov et al., 2012; Johnson et al., 2007) before being imported into Brainsight.

**Figure 3:**
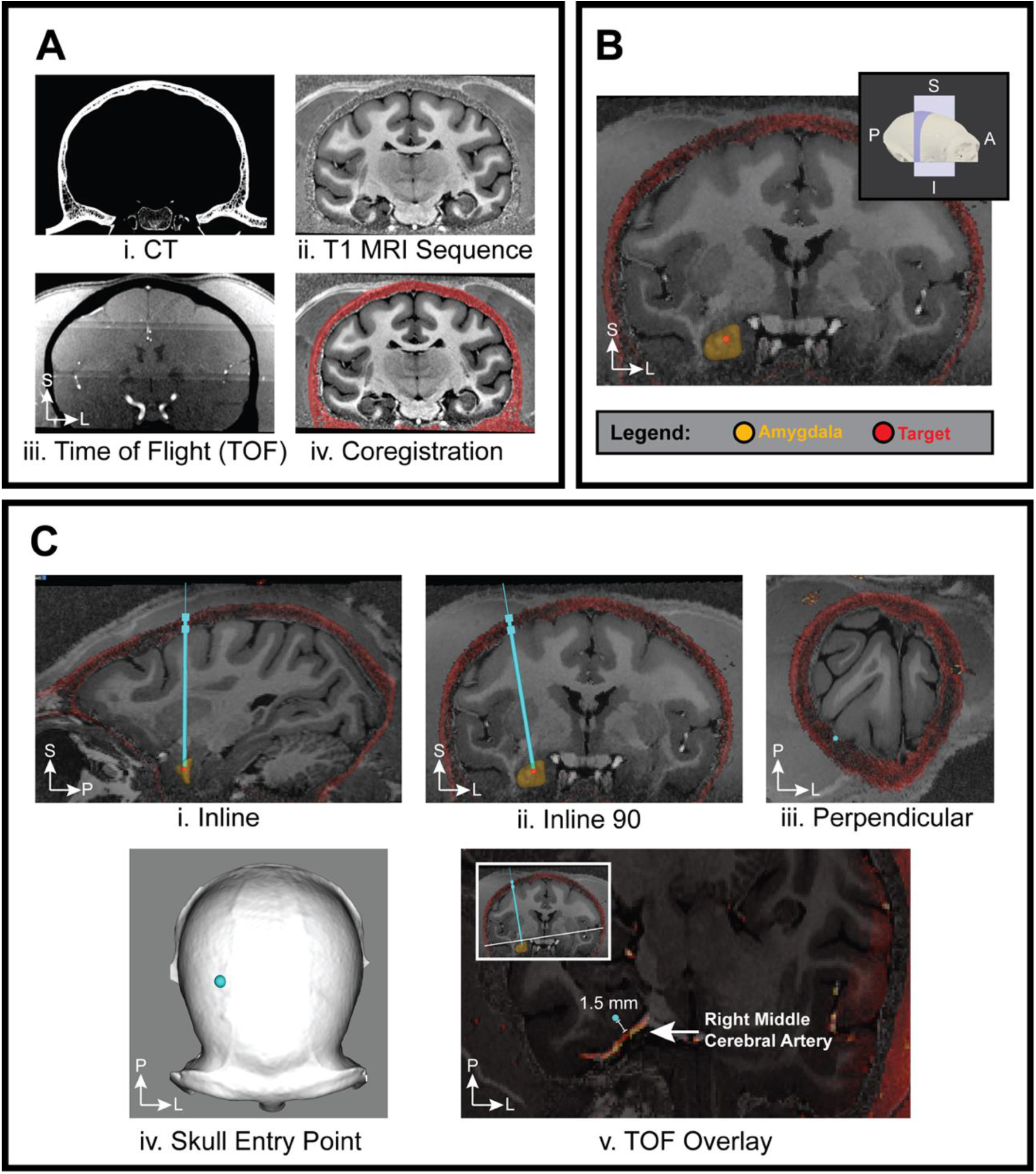
An overview of the steps to determine the trajectory of electrode implantation in macaque subject Mi. **A.** Image acquisition and co-registration, CT (dark red), MRI (gray scale). **B.** Visualization of CT, MRI, right amygdala reconstruction (yellow) and selected target (red) in Brainsight software. **C.** Selected trajectory (cyan) visualized in various views within Brainsight software. Views (i) - (iii) are in-line with the selected trajectory. (iv) Entry point of selected trajectory on skull reconstruction. (v) Verifying that major cerebral vasculature is avoided through overlaying trajectory on TOF sequence. Selected trajectory (cyan circle) is 1.5 mm from the middle cerebral artery. *Insets:* B. The illustrated coronal plane on the subject’s skull. C. Transverse slice, in-line of trajectory, in the plane of the middle cerebral artery. S = Superior; I = Inferior; A = Anterior; P = Posterior; L = Left; R = Right.

In Brainsight, the right amygdalae nuclei (yellow) were identified using a DTI atlas (Calabrese et al., 2015) with an interest in the basal, lateral, medial and central nuclei (Fig. 3B). After this region of interest (ROI) was defined and reconstructed (orange), the targeted nucleus, the basolateral amygdalae (BLA, red), was marked in the software and referenced for all subsequent targeting steps (Fig. 3B).

In determining viable trajectories to reach the target, a few considerations were made. To minimize the drift of the drill during a craniotomy, trajectories were optimized to be orthogonal to the skull surface. This was evaluated using a 3D skull reconstruction (Fig. 3C). The presence of major cerebral vasculature in proximity of the trajectory and beneath the skull-entry point was also taken into consideration. To evaluate this, the TOF sequence was superimposed on the MRI and the potential trajectory. We ensured that there was at least 1.5 mm of clearance from major vasculature such as the internal carotid or the middle cerebral artery (Fig. 3C).

In keeping with these considerations, three viable trajectories for targeting the right BLA were determined. Our preferred trajectory (cyan) is presented in Figure 3C.

Based on the chosen implantation trajectories, a cranial cap was designed and fabricated. Our lab has previously produced custom cranial caps for neurophysiological experimentation with NHP subjects, and the process from design to fabrication has been thoroughly documented (Blonde et al., 2018). While the design process is similar for each cap, the implantation protocol and the region of interest can influence each cap’s feature sets. The design for the current implants included a wide skull-access window, a bowl enclosure, and a designated region for the mounting of a headpost (Fig. 5). The window provided flexibility for performing multiple implants, while the bowl enclosure allowed protection of the implanted assembly and housing for the MBA connectors. The headpost was used for head fixation to collect eye data during performance of behavioural paradigms. While the figures in this manuscript present the cranial cap design & cap implantation process for monkey Mi, a similar protocol was followed for monkey Ma’s cranial cap.

As the design was developed and refined, it was important to verify its fit and function with the trajectory and the tools used. This was achieved through both software and hardware confirmation.

#### Software Confirmation

First, the cranial cap prototype was imported into Brainsight. It was overlaid with the 3D skull reconstruction to confirm that the trajectory lies within the skull-access window and that there was sufficient margin for the anchor bolt to be fastened (Fig. 4A). Using the length of the MBA, we confirmed its fit within the bowl enclosure in software as well (Fig. 4B).

**Figure 4:**
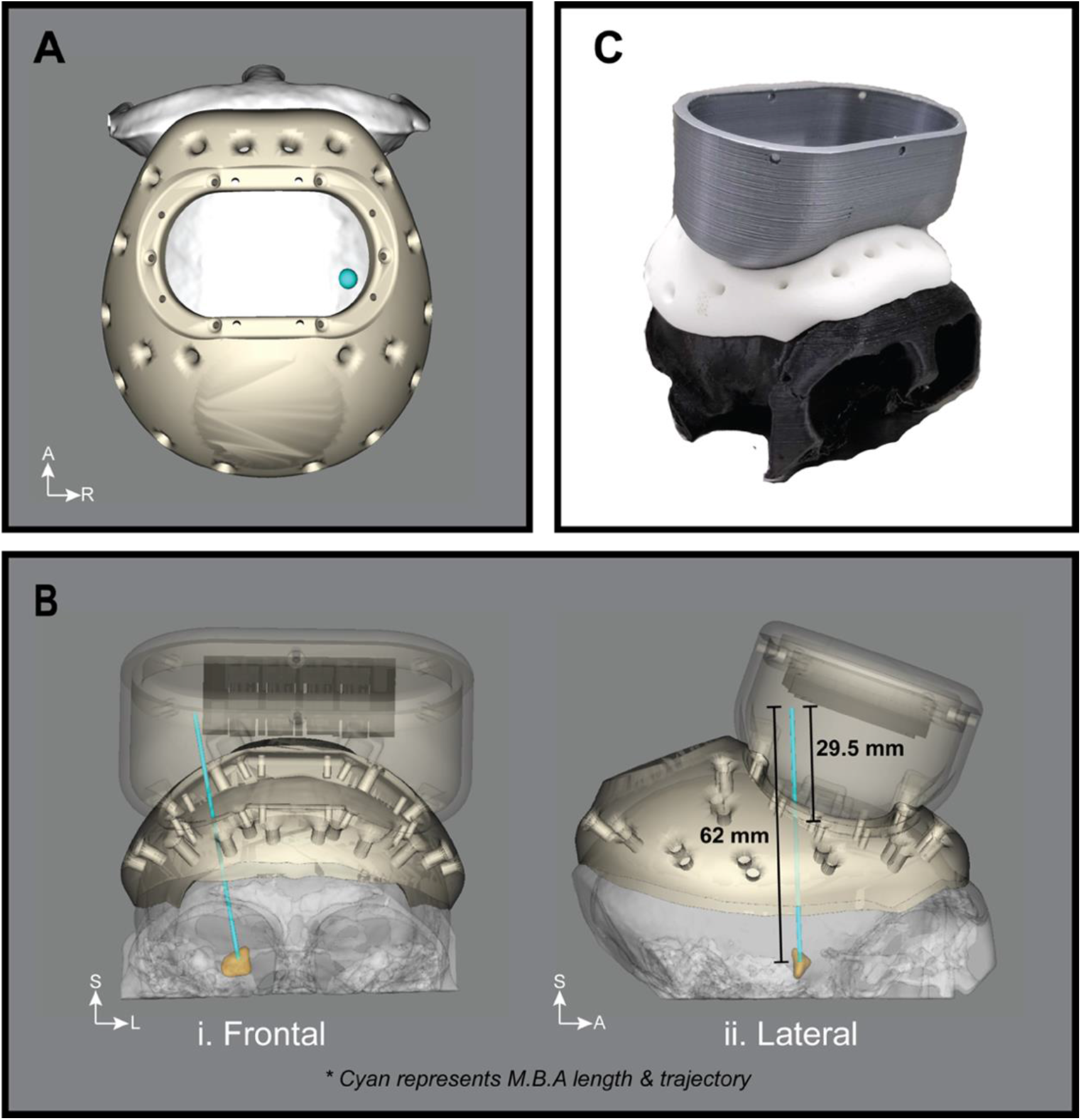
Confirmation of cranial cap and trajectory compatibility through software and hardware techniques during the cap design and development process (Monkey Mi). **A.** Cranial cap’s skull-access window margin accommodated the trajectory and the anchor bolt. The diameter of the cyan sphere reflects the diameter of the anchor bolt. **B.** A bowl enclosure was used to house the assembly and connectors. Measurement of assembly length, in-line with trajectory to the target (orange) within software to ensure fit inside the bowl. **C.** 3D-printed prototype of cranial cap and bowl to test fit with skull contour, cap screw accessibility and mock surgery procedures to test assembly fit. S = Superior; I = Inferior; A = Anterior; P = Posterior; L = Left; R = Right.

#### Hardware Confirmation

At various points in the design process, a 3D prototype was printed in polylactic acid (PLA) and nylon to validate the functionality of the cranial cap (Fig. 4C). This allowed us to address issues of cap fit with the skull contour and whether all screw holes were easily accessible using available drivers. Using these prototypes, we also performed a mock surgery to execute the implantation protocol using the final cranial cap design and ensuring that all hardware components could be housed within the bowl enclosure. These validation techniques allowed us to iterate upon our cap design until finalized. The cranial cap was machined in PEEK (Polyether ether ketone) for monkey Mi (Fig. 5) and 3D printed in titanium for monkey Ma for implantation.

**Figure 5:**
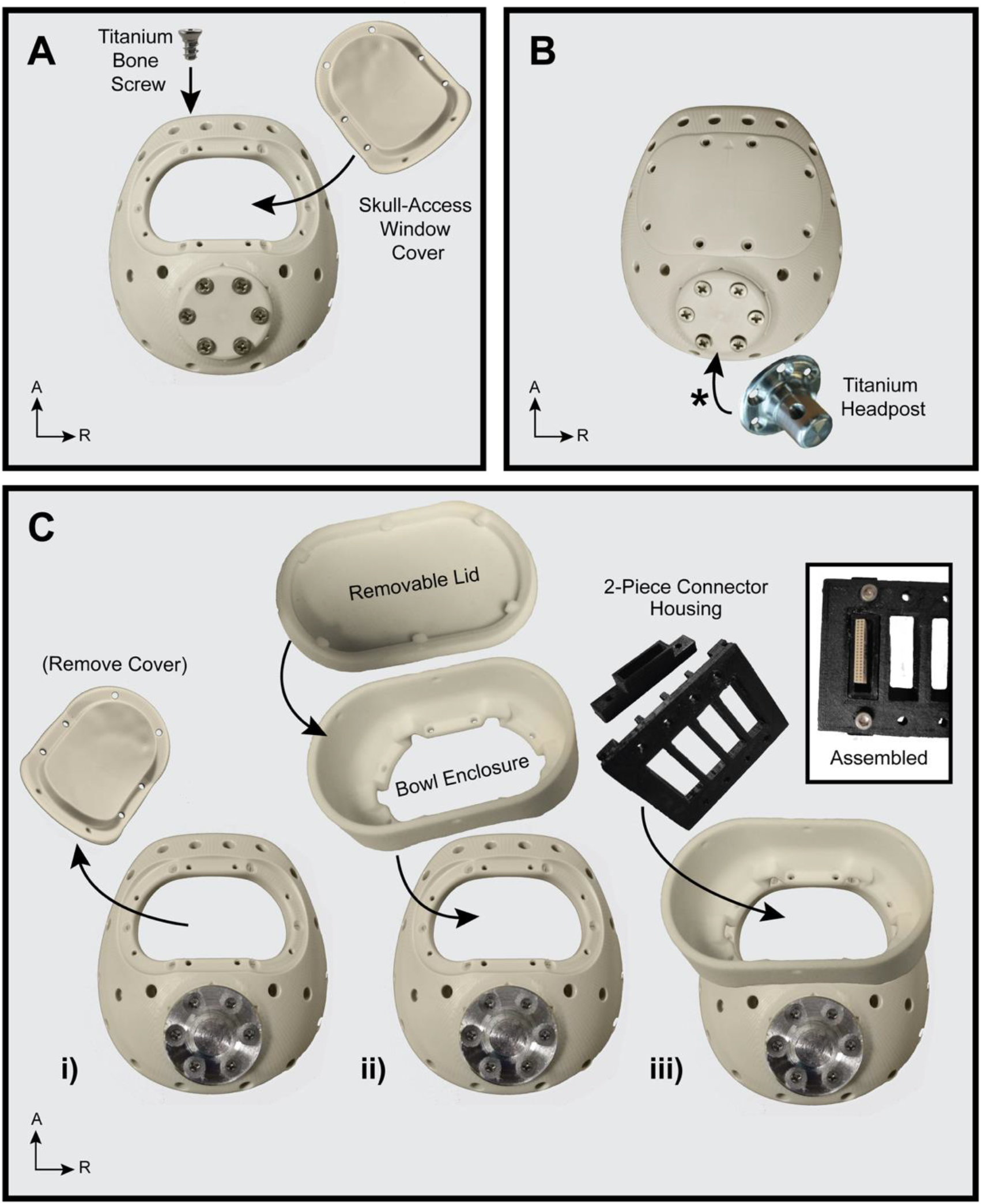
Overview of cranial cap features and components for monkey Mi at various stages of implantation. **A.** Reflects the elements used for the initial cranial cap implant. Window is attached to the cap, which is implanted on the skull using titanium bone screws. The headpost mounting base (posterior) is covered. **B.** A titanium headpost will be attached to the designated mounting base, after removal of the cover*. **C.** Cap components involved in electrode implantation. (i) Window cover is removed and (ii) replaced with the bowl enclosure and lid. (iii) Connectors will be secured inside the bowl using a 3D-printed two-piece housing (black).

### Cranial cap components

Before the initial cranial cap implantation, the skull-access window was closed using a cover fastened to the cap with screws (Fig. 5A). The cap with the attached cover was fixed to the skull using titanium bone screws (Gray Matter Research, Bozeman, MT). The next stage was the implantation of the titanium headpost (Fig. 5B). This was mounted to the designated base on the posterior region of the cranial cap. This base included a cover plate to ensure a smooth surface to avoid irritation of the overlaying skin post cap implantation and was subsequently removed before mounting of the headpost in a separate procedure.

The final stage reflects the components that were used during and after the electrode implantation (Fig. 5C). First, the window cover was removed and replaced with the nylon-printed bowl enclosure used to house the implanted assembly. This was covered with a removable lid, allowing access to connectors for neural recording. This bowl enclosure was fixed directly to the cap using screws.

A custom two-piece connector housing was 3D printed using PLA to secure the electrode connectors and provide easy access within the recording set-up when attaching the headstage (Fig. 5C). Each 32-channel connector was embedded into a smaller connector holder using dental cement. This holder was then screwed onto the connector bridge, which held up to four connector holders (Figure 5C iii). This bridge was then secured to the walls of the bowl enclosure using dental cement. Should another electrode implant be planned, each connector holder could be detached from the bridge, allowing the bridge to be removed along with the bowl enclosure, revealing access to the skull.

### Custom 3D Printed Electrode Microdrive

The microdrive parts consisted of a drive screw which was soldered into a nut, a body that screws onto an anchor bolt, a two-piece electrode clamp, and a cover (Fig. 6A). It was designed using Autodesk Fusion 360 software and printed using Formlabs 3B resin 3D printer (Appendix A).

**Figure 6:**
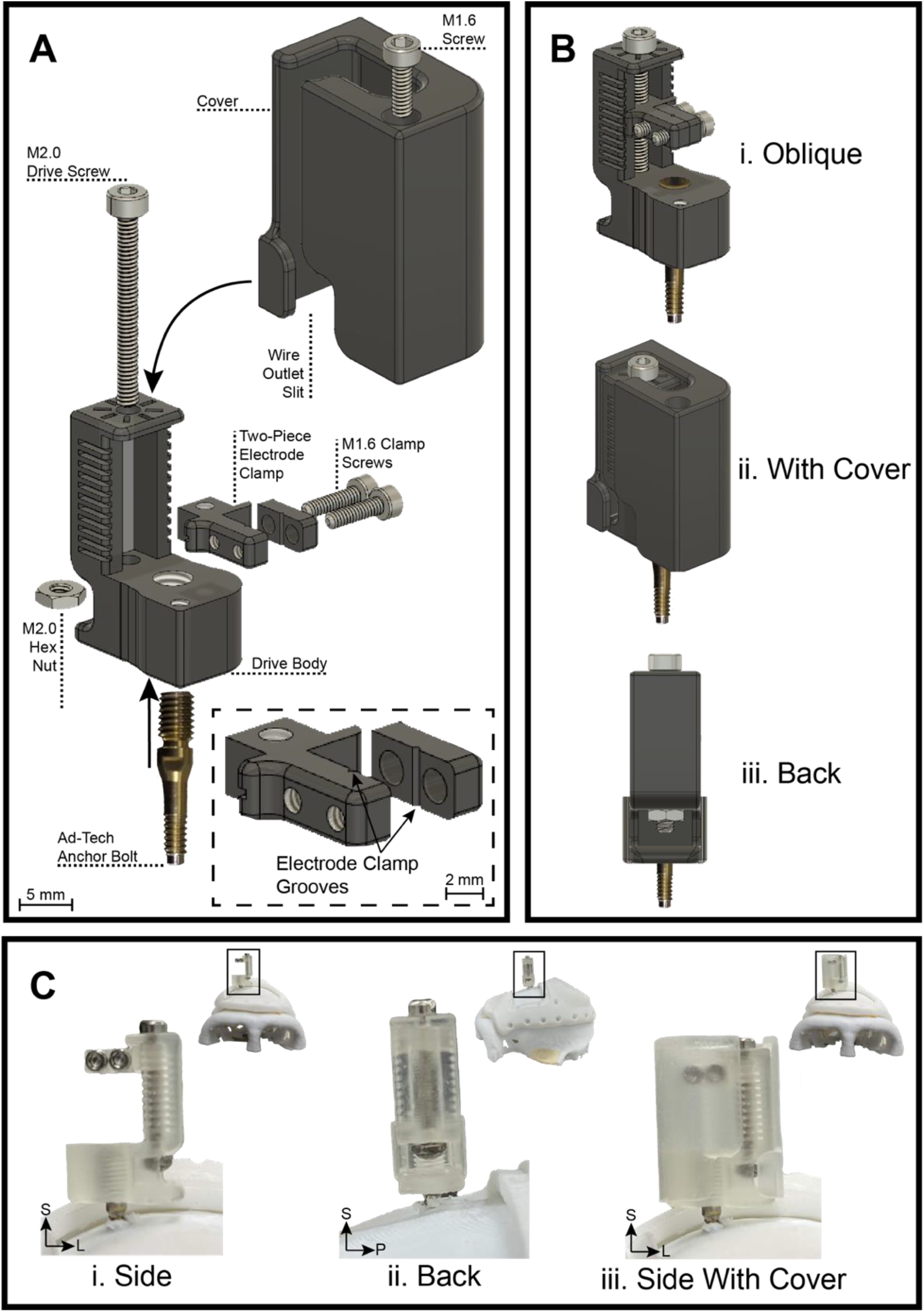
Schematic and implementation of custom 3D-printed microdrive which allowed single-axis movement of up to 10 mm. **A.** Blow-out schematic of microdrive components. The M2.0 drive screw is passed through the top of the drive body and screwed through the two-piece electrode clamp. The screw is held in place by the M2.0 hex nut, which is soldered (see C-ii). The electrode is held within the electrode clamp grooves by two M1.6 clamp screws. The microdrive is screwed onto the Ad-Tech anchor bolt, fixing it in-line with the desired trajectory. A cover, screwed onto the drive body, was designed to protect the microdrive and electrode from the fluids that can build up inside the bowl enclosure. It also contained an opening along the sidewall as an outlet for the connector wires and ground during insertion. **B.** Assembled schematic views of the microdrive. **C.** Views of 3-D printed microdrive. The microdrive implant is oriented in the medial-lateral axis to accommodate it within the cap. In (i) and (ii), the soldered nut is visible. This allowed the drive screw to turn in place for the clamp to move along a single axis. (iii) Two mirrored versions of the cover were made, with the wire outlet slit being on either side. One is shown in schematic B(ii), and the other in C(iii). The version used was dependent on the microdrive orientation during implantation. *Insets:* (i) Frontal view of mock subject, post-implant; (ii) Side view of mock subject. (iii) Frontal view of subject. S = Superior; I = Inferior; A = Anterior; P = Posterior; L = Left; R = Right.

The electrode was secured in between the grooves of the two-piece clamp by tightening the clamp screws (Fig. 6A). The clamp was held in position by the drive body and drive screw. Turning the drive screw rotated the screw in-place (enclosed by the body and the soldered nut below) and allowed raising or lowering of the clamp. The microdrive was positioned in-line with the desired trajectory by threading it onto an anchor bolt already secured on the skull. The cover was placed over the microdrive assembly, top-down, and secured onto the drive body using a screw. Figure 6B (3-D schematic) and Figure 6C (Hardware Prototype) show the completed assembly of the microdrive. All hardware parts shown in Figure 6 are available and referenced to the manufacturer website in supplementary table 1. Microdrive CAD files (STEP and STL files) as well as drawings of the parts are made available on an online repository (https://doi.org/10.5281/zenodo.6877605).

### Electrode assembly within bowl enclosure (schematic)

Figure 7 summarizes our protocol for the procedure within the cranial cap bowl. The anchor bolt, guide tube and electrode assembly were secured in-line with the trajectory. Dental cement was applied at the MBA-guide tube-anchor bolt interface securing the components as a unit. The guide tube was positioned above the target, allowing the microwires to splay as the MBA exits and travels towards the target. Finally, the entire assembly was housed within the bowl enclosure of the cranial cap. The MBA connectors were secured into the connector holder using dental cement and attached to the connector bridge using screws. The entire connector bridge was then attached to the wall of the bowl enclosure using dental cement.

**Figure 7:**
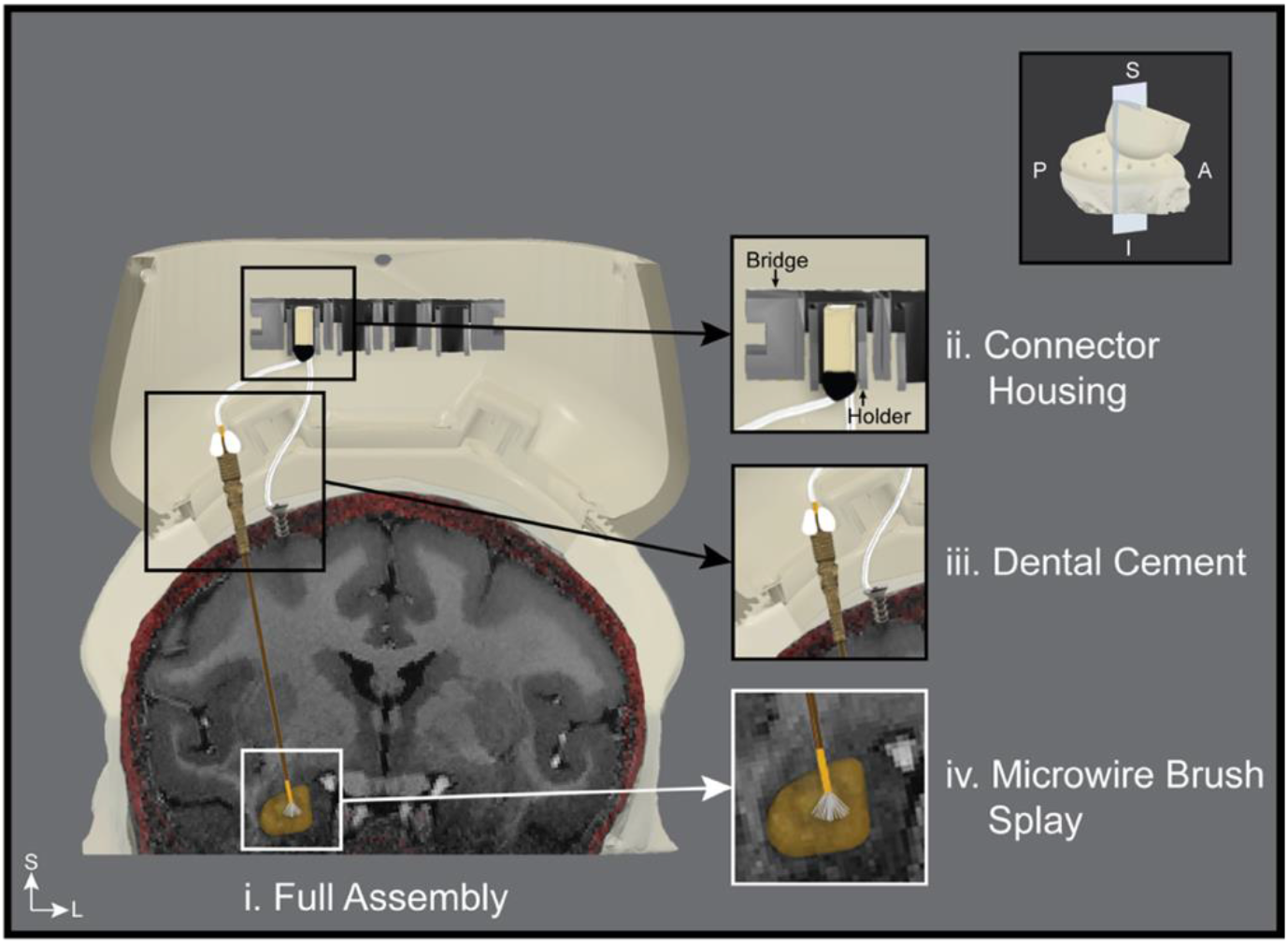
Demonstration of the full electrode assembly within the cap. Inset: coronal section of the subject skull with cranial cap in the plane of the trajectory. (i) An overview of the assembly, in-line of trajectory, and its placement within the cap housing. (ii) The MBA connector is fixed into the holder using dental cement. The holder with the connector is attached to the bridge using screws. The bridge* is attached to the walls of the bowl using dental cement. (iii) Dental cement at anchor bolt base (not shown) and proximal opening. (iv) Splaying of microwire brush array within the target after exiting the guide tube. Note*: The orientation of the connector bridge in this schematic is for demonstration purposes only. The true orientation is rotated by 90 degrees. S = Superior; I = Inferior; A = Anterior; P = Posterior; L = Left; R = Right.

### Cranial Implant Surgeries

#### Cap Surgery

To implant the cranial cap, a C-shape incision was made for the scalp and the temporal muscles to be retracted (Fig. 8A). The skin was kept hydrated while the skull surface was cleaned with a bone scraper to position the cranial cap flush with the skull. As mentioned earlier, the skull-access window cover was attached to the cap before the procedure. Titanium bone screws were used to secure the cap to the skull. Once the screws were in place, the skin and muscle tissue were placed back over the implanted cap and the skin was sutured.

**Figure 8:**
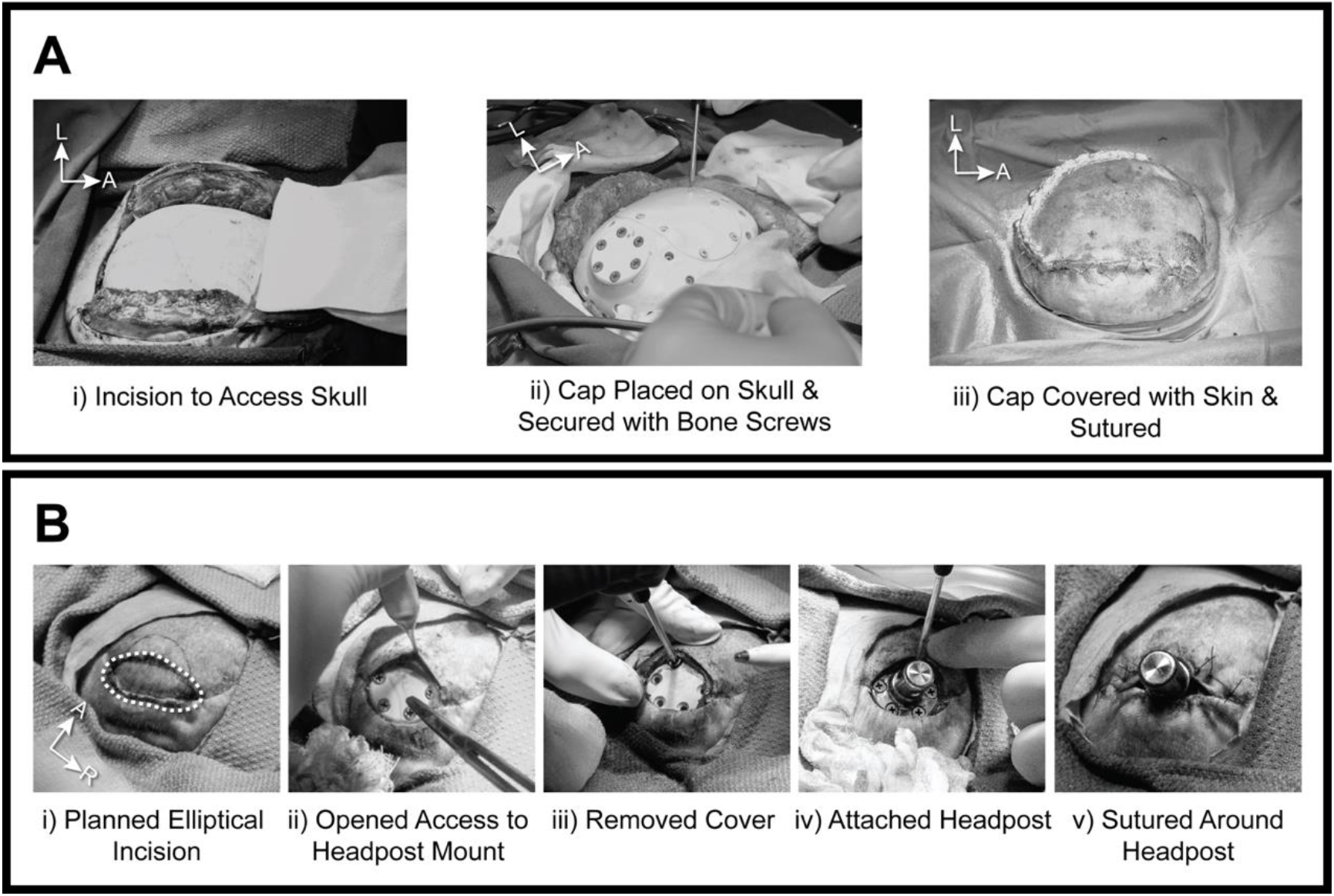
Cranial cap and headpost surgical procedures (monkey Mi). **A.** Cranial cap surgery. (i) Skin and muscle tissues are retracted. (ii) Cranial cap was placed on the skull and cranial screws fastened it to the bone. (iii) Skin and muscle are pulled over the cap and sutured. **B.** Headpost mounting surgery. (i) Dotted line indicates where the incision was performed to access the headpost mounting base (ii) on cranial cap. (iii) Remove headpost cover plate before (iv) attaching headpost. (v) The skin was sutured around the headpost. S = Superior; I = Inferior; A = Anterior; P = Posterior; L = Left; R = Right.

#### Headpost Surgery

Approximately 4-6 weeks after the cap surgery, the headpost-mounting surgery took place (Fig. 8B). First, an elliptical incision was made in the posterior region of the cap, crossing the boundary of, while avoiding the initial surgical scar. This allowed access to the mounting base of the headpost on the cranial cap. The incision was opened and the cover plate on the base was unscrewed and removed. The titanium headpost was then secured to the cranial cap using screws. The skin was sutured around the headpost, allowing access to the post for head fixation in the primate chair (Fig. 8B v).

Approximately 3 weeks after the headpost surgery, when head fixation training was completed, the subject underwent a procedure to remove the skin covering the top of the cranial cap. This is to provide access to the implantation window and screw fiducials described in the next section.

### Fiducial marker determination

Following the cap and headpost surgeries, a new CT scan was obtained and co-registered within the 3D Slicer software with the MRI-aligned pre-cap CT. The Markups Module was used to specify anatomical point pairs on each CT scans, and the Fiducial Registration Module was used to perform a rigid co-registration between the indicated point pairs. Post-cap implantation CT scan allowed for visualization of cranial cap screws, which present themselves as hyperintensities on the CT (Fig. 9A i) and can be easily isolated. Figure 9A(ii) presents screw reconstructions in Monkey Mi’s implant that was produced in Brainsight software through intensity thresholding. These reconstructions were then overlaid with the post-cap CT skull reconstruction and a cranial cap model, as the implanted PEEK cranial cap does not present itself well on the CT. Cap screws were used as fiducial markers for registration. In neuronavigation, registration fiducials are salient features identified on anatomical images (e.g. CT scan) within a surgical planning software pre-operatively, and are physically localized during surgery to achieve subject-image registration. This allows for real-time tracking of surgical instruments within the anatomical scans in the Brainsight software during surgical operation, to locate pre-determined electrode implantation trajectories.

**Figure 9:**
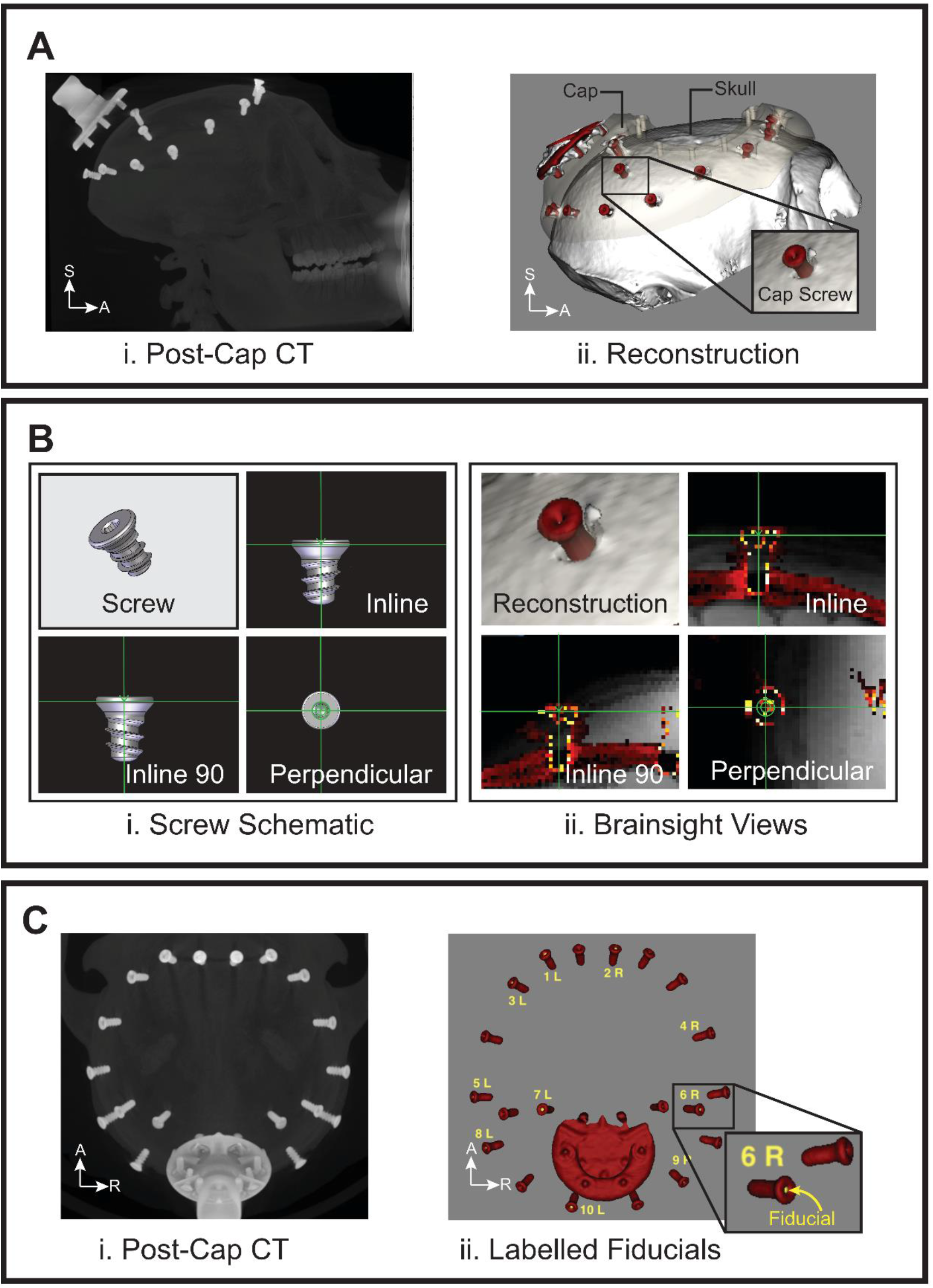
Determining physically accessible fiducials in Brainsight software pre-operatively for subject-scan registration during the electrode implantation surgery. **A.** (i) Post-cap implantation CT is used to produce a (ii) skull reconstruction (white) with cranial cap (beige) and screws (red). **B.** Software fiducials (yellow; visible in inset in C) will be placed inside 10 selected cranial cap screw drives (i - schematic; ii - Brainsight). Graphic image of cranial screws provided by Gray Matter Research and modified with permission. **C.** (i) Top-down view of post-cap CT, presenting (ii) screws containing labeled fiducials (yellow) to be referenced during surgery. S = Superior; I = Inferior; A = Anterior; P = Posterior; L = Left; R = Right.

Cranial cap screws were used for the placement of these markers as they are stable, easily accessible, and could be consistently located. In software, fiducials (Fig. 9C ii, yellow) were placed within the drive of the screw, in orthogonal planes as seen in different views of Figure 9B. The orientation of the fiducials determines how they need to be localized for an accurate registration. Ten screw fiducials were placed asymmetrically around the target as the centroid and were labelled for ease of referencing during surgery.

### Surgical equipment

#### Individual hardware

Surgical apparatus included neuronavigation (software and hardware) and implantation equipment from Brainsight (Rogue Research Inc., Montréal) and Ad-Tech Medical Instruments (Oak Creek, WI). This consisted of equipment for positioning the subject (Fig. 10A), head fixation (Fig. 10C), neuronavigation hardware (Fig. 10B) and equipment for implantation of the electrode (Fig. 10D). The surgical chair allowed intubation of the subject in supine position while the C-clamp attachment (Fig. 10C) fixed the head using four skull pins (Fig. 10C ii, Fig. 11A,). To perform the surgery using neuronavigation, an infrared position sensor camera (*Polaris Vicra*, Fig 10B i) was used to locate trackers (Fig. 10B ii-iii) indicating the subject position and the 3D pose of the pointer. The pointer was used to locate the fiducials for registration of the physical space of the subject and anatomical images, and for matching predetermined trajectories.

**Figure 10:**
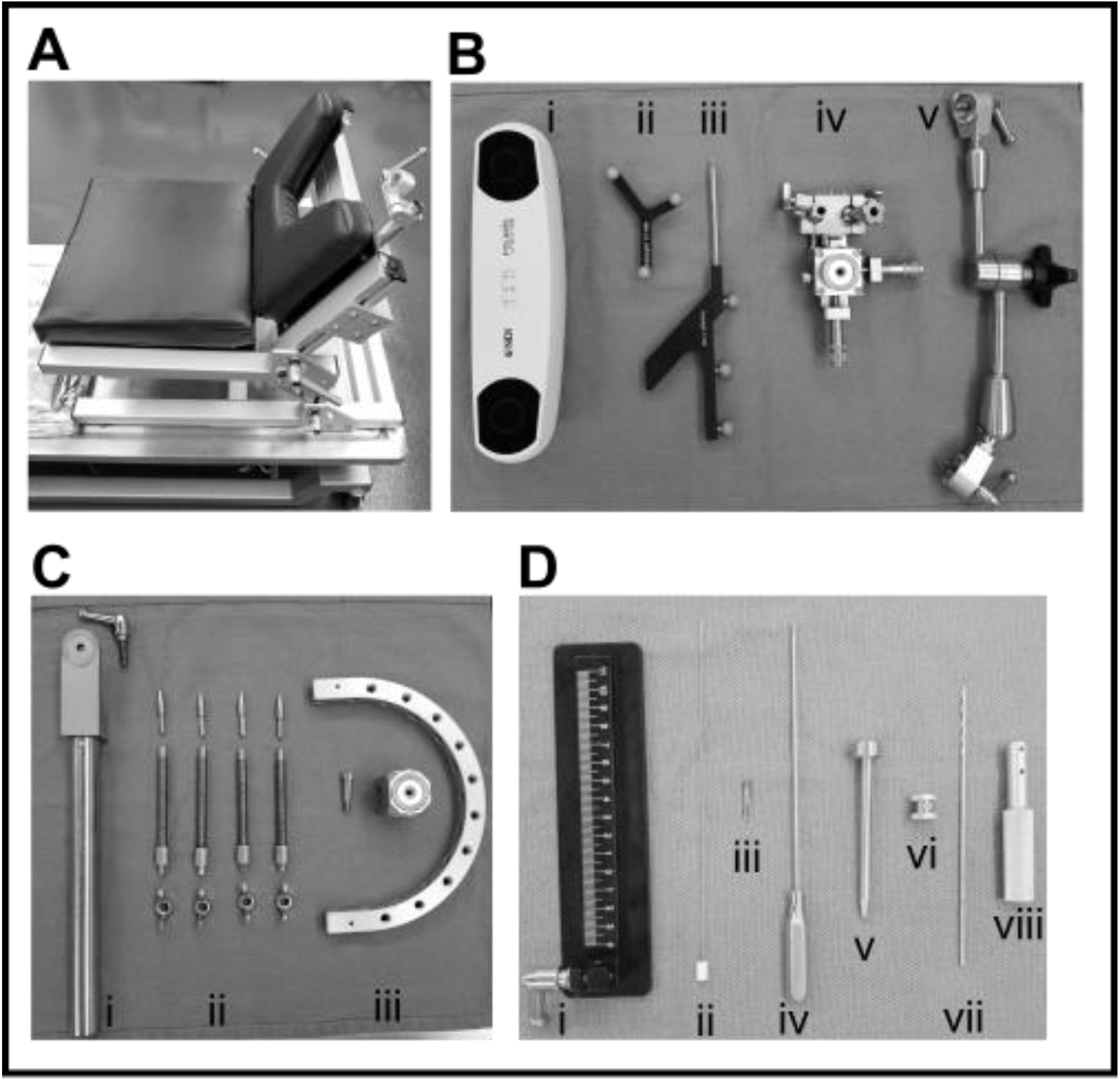
Individual hardware for subject positioning, neuronavigation, and implantation. **A.** Primate chair from Brainsight seated the subject in supine position at the desired angle. Joining adapter in the chair’s backrest allowed for connection of the fixation arm holding the C-clamp for head fixation (as shown in Fig 11C). **B.** Neuronavigation hardware: (i) Polaris infrared camera tracked reflective spheres on the (ii) subject and (iii) pointer trackers. iv) Freeguide (FG) arm stage which allowed mounting of equipment into two separate chucks. This stage was attached to the (v) FG arm. **C.** The C-clamp and attachments: (i) Fixation arm which connected to the joining adapter on the primate chair. (ii) Four sets of butterfly nuts, skull screws and pins were passed through the (iii) C-clamp holes which allowed for fixation of the subject head. Starburst block and attachment screw (iii) allowed for attachment of C-clamp to the fixation arm (i) and attachment of the FG arm. **D.** Implantation hardware: Ruler (i). Blunt stylet (ii) was inserted prior to the guide tube. The anchor bolt (iii) was fastened into the skull with the placement wrench (iv) through the drill sleeve (v). Drill bit stopper (vi) allowed the drill bit (vii) used along with the handle (viii) to stop at the appropriate depth.

**Figure 11:**
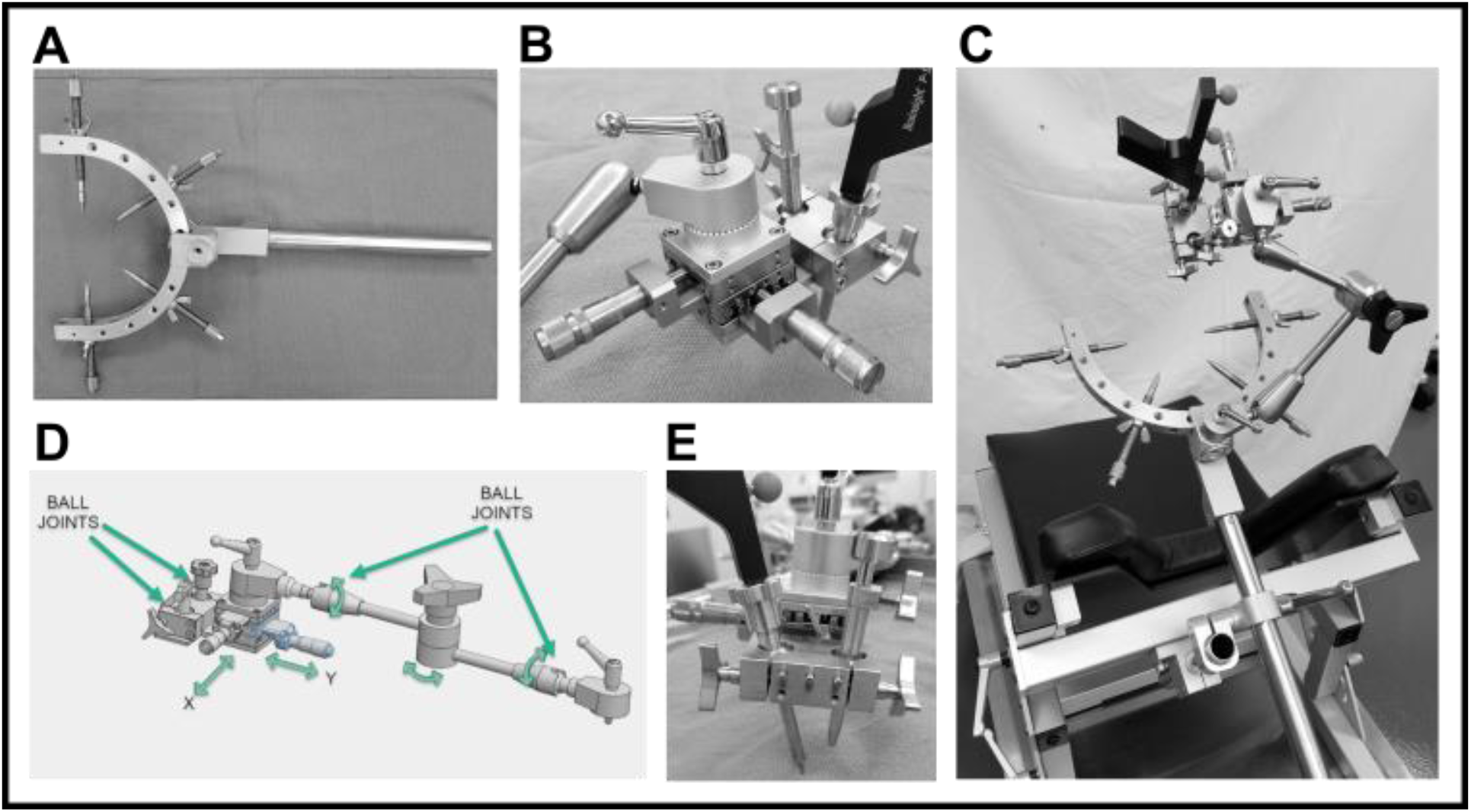
Assembled Hardware Components. **A.** Assembled C-clamp. Included are all components in Figure 10C. **B.** Close up of the FG stage with the pointer and drill sleeve inserted into the chucks of the stage. **C.** The complete assembly: The C-clamp system attached to the primate chair with the FG arm, stage, and the inserted pointer and drill sleeve. **D.** Schematic of the anatomy of the FG arm and stage. Joints along the arm allow high degrees of freedom for rotation of the arm and stage. The x-y adjustments on the stage enable translation of the mounted tools, while the ball joints on the stage allow for rotation of the tools. Image courtesy of Rogue Research Inc. **E.** Pointer and drill sleeve inserted in the stage chuck. Knobs on the side of the stage loosen the ball joints, allowing for rotation of the mounted tools.

#### Assembled hardware

The Freeguide (FG) arm (Fig. 10B v, Fig. 11D), along with the attached stage (Fig. 10B iv), allowed for stabilized movement of the pointer during trajectory mapping. The stage chuck permitted mounting of the pointer and surgical instruments (Fig. 11B, C, E) during implantation, with the ball joints allowing rotation of the equipment on the stage (Fig. 11D). Lastly, the *x-y* translation of the stage allowed for fine-tuning of the trajectory (Fig. 11D) after locking the position of the FG arm in space. Figure 11C illustrates the complete assembly of the apparatus attached to the C-clamp.

### Electrode Implantation Surgery Overview

An overview of the surgical protocol for both animals is presented in supplementary Figure 2, with animal specific protocols highlighted. In summary, the animal was first positioned and its head fixated. Using the Brainsight system, the subject and the anatomical images which were used for trajectory planning pre-operatively were then registered. This allowed the use of neuronavigation in locating pre-operatively determined trajectories, and the electrode insertion protocol was then carried out.

### Subject positioning

The subject was positioned on the Brainsight chair within the range of the Polaris camera, with the Brainsight computer containing the surgical plan being operated by an assistant (Fig. 12A). This computer was used for registration of the subject as well as visualizing the 3D pose of the pointer during mapping of the trajectories in real-time. The subject was seated in a supine position and small incisions were made in the temporalis muscles to reach the bone in four locations below the cap perimeter. These incisions allowed for insertion of four skull pins, which stabilized the head within the C-clamp frame (Fig. 12B). Once the subject’s position was finalized, the surgical area was draped, and the bowl enclosure was disinfected with an iodine solution and removed to provide access to the implantation window (Fig. 12C). The exposed skull surface was disinfected and cleaned to gain access to the bone. The subject tracker was then secured to the C-clamp (Fig. 12D), ensuring that the orientation of the reflective spheres was optimized for detection by the Polaris camera seen in Figure 12A.

**Figure 12:**
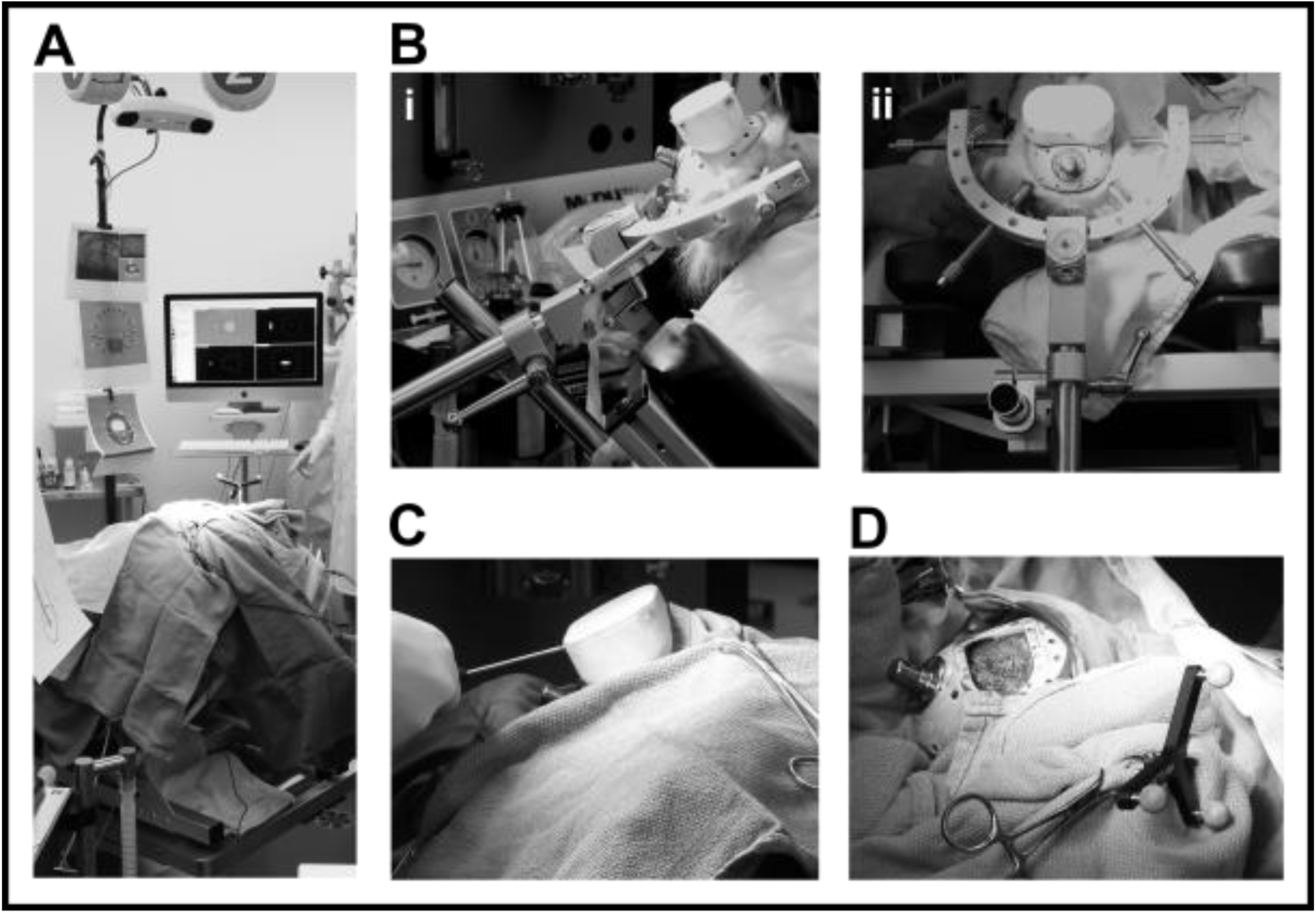
Subject positioning and subject tracker placement. **A.** Finalized surgical set up with the subject head fixed on the chair and draped. Polaris camera positioned for optimal visibility of the subject tracker. Surgical assistant operating the Brainsight computer. **B.** Subject in supine position, head fixated using the C-clamp system shown from the lateral (i) and posterior (ii) views. **C.** Bowl enclosure removal for access to the skull. **D.** Once the skull surface was cleaned, the subject tracker was attached to the C-clamp and its visibility by Polaris camera was verified.

### Neuronavigation

#### Subject-Image registration

To locate the predetermined trajectories from Brainsight software in physical space, registration of subject (physical) space with anatomical image (software) space is required. Homologous fiducials on the subject’s head and images were identified and matched, transforming the subject space into “image” space (Arun et al., 1987). Matching these coordinate systems allows for the 3D pose of tracked tools (e.g. pointer) to be represented in the anatomical images, enabling real-time navigation through the scans. To begin fiducial-based registration, the Polaris camera was positioned to capture both the fixed subject tracker and the pointer’s range of movement (Fig. 13A-B). Once a fiducial was selected in the software, the surgeon positioned the tracked pointer orthogonal to the head of the screw and the tip was rested in the screw drive (Fig. 13C). The fiducials had been placed in the cavity below the drive surface and could consistently be located by the pointer’s tip (Fig. 13C inset). Once the pointer was placed on the presumed fiducial location, its tip position was sampled by the Brainsight system to register the physical and software fiducial. This step was repeated for all ten fiducials (Fig. 13D i). To validate the registration, the pointer was positioned at various locations on the cap. The discrepancy between the position of the pointer tip in physical space and in the anatomical images was observed and corrected by repeating the registration process (Fig. 13D ii-iv). A view of the Brainsight software during the verification process is shown in Supplemental Figure 1.

**Figure 13:**
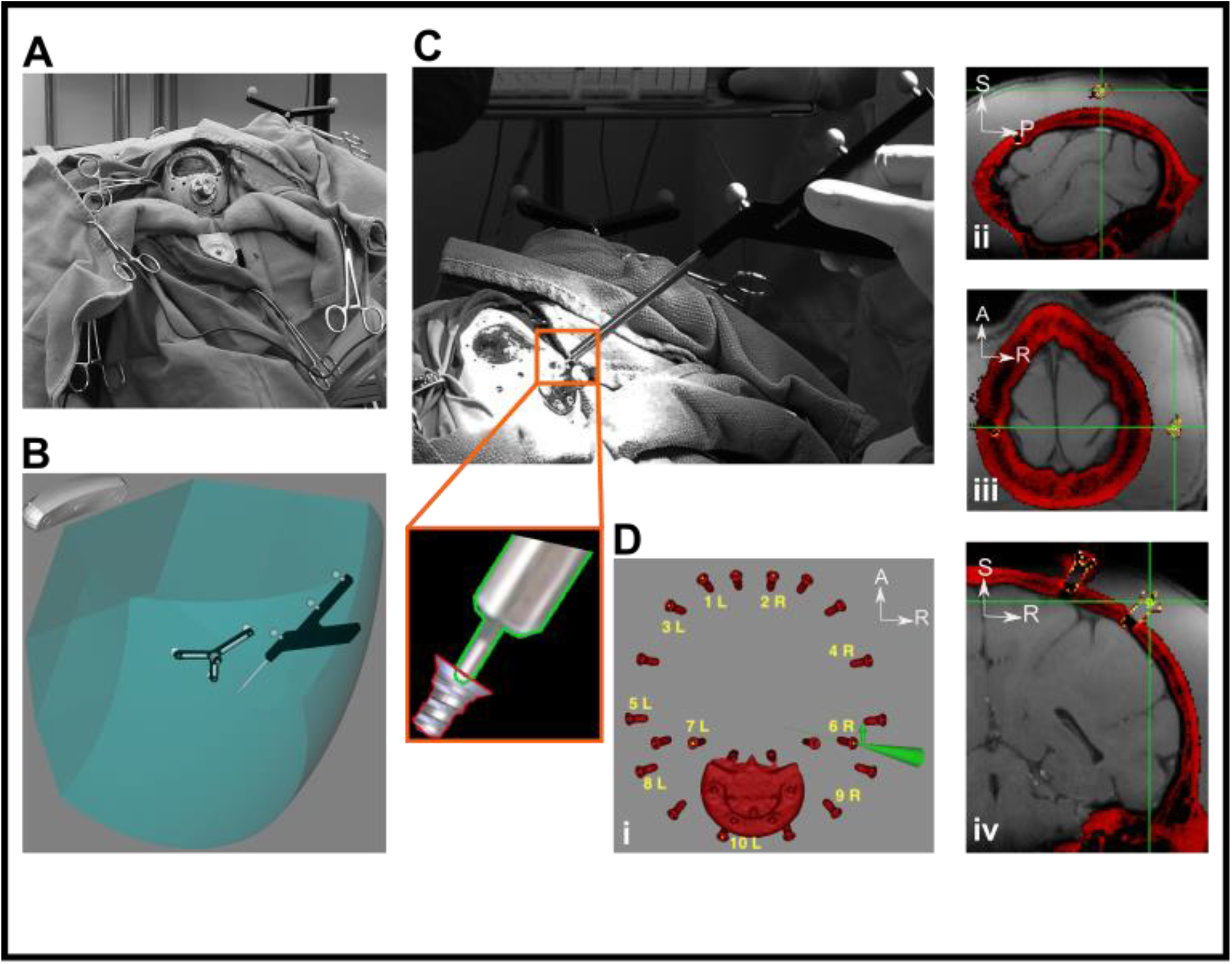
Subject-Image registration. **A.** Subject positioned in supine position with the subject tracker firmly fixed to the C-clamp. **B.** Verification of the position of the Polaris camera to capture both the subject tracker and pointer. **C.** Sampling of fiducial markers. Pointer is rested on the chosen fiducial (cranial screw) by the surgeon. Inset shows a lateral view of the position of the pointer’s tip in relation to the screw drive. Fiducials were placed below the screw drive in the software and located physically during the registration. Graphic image of cranial screws provided by Gray Matter Research and modified with permission. **D.** Top view (i) of the cap screw reconstructions and the ten screws that were used for placement of fiducials (yellow) for registration. Green cone represents the pointer tracked in software. The validity of the registration was assessed by placement of pointer tip at various points on the cap. (ii)-(iv) shows placement of pointer (green cross) onto one of the co-registered fiducials (yellow) in sagittal, transverse, and coronal views respectively. S = Superior; I = Inferior; A = Anterior; P = Posterior; L = Left; R = Right.

#### Matching implantation trajectory

Registration of the subject space with the anatomical images allowed for real-time navigation to locate the pre-planned electrode trajectory. A few trajectories were planned to allow for flexibility during the electrode implantation surgery. Once a trajectory was selected in the software (Fig. 14A), the tracked pointer was used to find the chosen trajectory in physical space. To do this, the FG arm was first attached to the C-clamp. The stage of the FG arm allowed for mounting of the equipment, such as the pointer (Fig. 14B). The FG arm allowed for stabilized but extensive movement of the pointer around the subject’s head for locating the pre-planned trajectory. The joints of the FG arm (orange arrow heads, Fig. 14B) allowed for 6 degrees-of-freedom of movement in 3D space (rotation and translation), while fastening them secured the arm in place. The steps for finding the planned trajectory were as follows: 1) The FG arm joints were loosened to allow for flexible movement with both hands to find the approximate angle of entrance (Fig. 14C). Feedback information about the error of the trajectory was communicated through the Brainsight system (Fig. 14B-E Insets). 2) Once the entry point and the approximate entry angle was found, the arm was locked in space and the ball joint on the FG arm stage (Figure 11D) allowed for adjustment of the entrance angle by permitting a cone of rotation (visualized as an orange cone in Fig. 14D). 3) As the trajectory angle was finalized, the x-y adjustment controls of the stage allowed for fine-tuning of the entrance point by translation of the trajectory in the x-y axes (as seen by x-y arrows in Figure 11D). 4) The finalized trajectory was then confirmed in the software and any deviation from the intended trajectory and target was reviewed (Fig. 14E).

**Figure 14:**
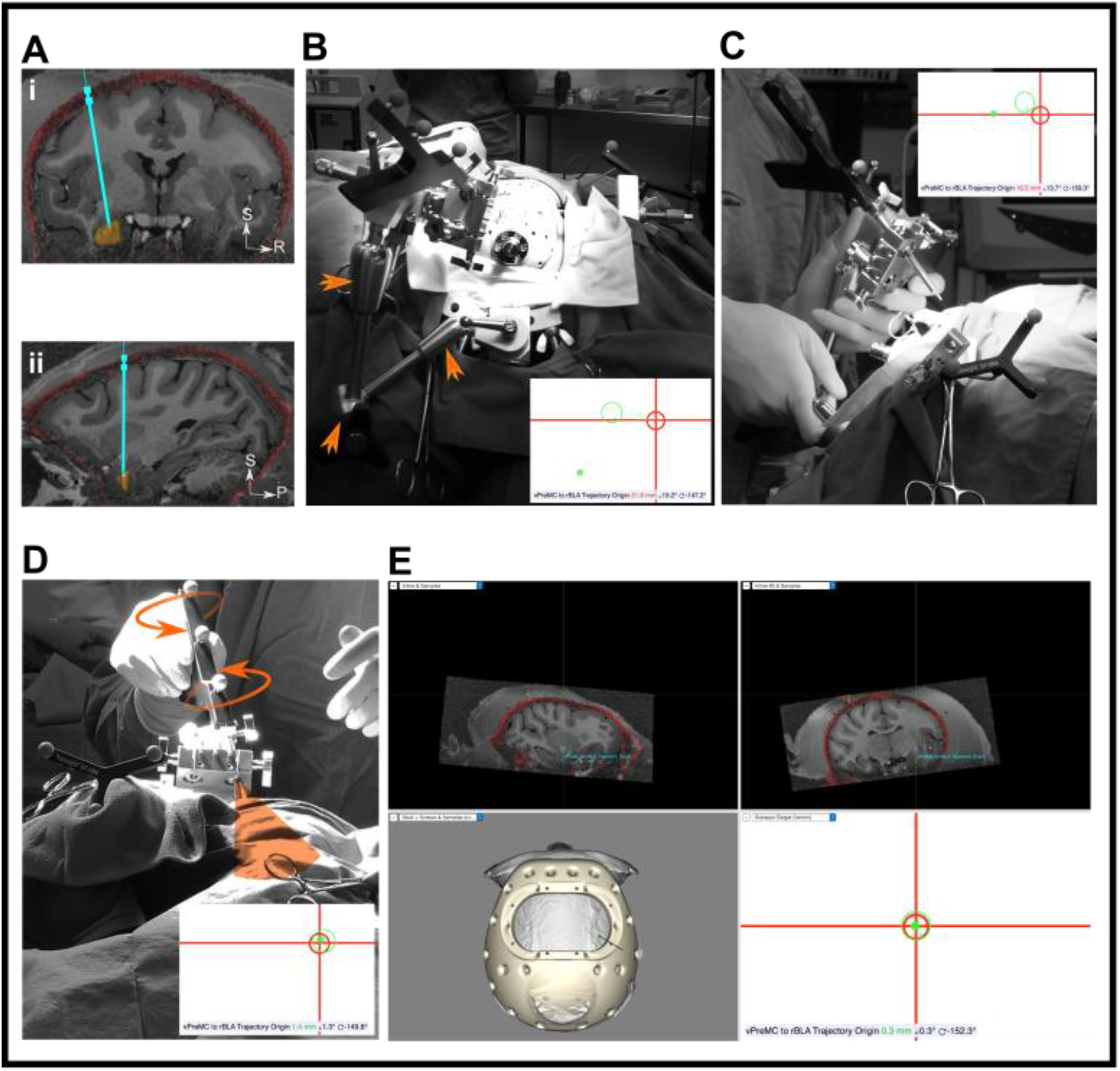
Neuronavigation steps. **A.** Predetermined trajectory targeting basolateral amygdala was selected. **B.** The FG arm was attached to the C-clamp and the pointer was mounted into the stage chuck to allow for stabilized navigation with the arm for finding the trajectory. Orange arrows indicate joints of rotation allowing for 6-degrees-of-freedom of movement of the FG arm. **C.** Both hands were used to move the FG arm to approximate the angle and entry point of the trajectory. **D.** Once the arm was locked, the ball joint on the stage was loosened allowing for rotation of tools mounted into the chuck. Shown here is the pointer rotation using the ball joint (possible rotation direction shown by orange arrows). This allowed for the range of rotation exemplified by the orange cone. **E.** Brainsight window after the trajectory was found. Two top panels show views from different sections, in-line with the pointer. Bottom left panel showing a top view of the skull and cap. Along with the insets, the bottom right panel shows the bulls-eye view of the error from the intended trajectory: Red Cross and circle represent the selected trajectory’s entry point and angle respectively. Green circle represents the angle of the pointer in space while the green dot represents the pointer tip (entry point). For successful targeting, the green dot (pointer tip) must be centered on the cross (trajectory entry point) and the green circle (pointer angle) aligned to the red circle (intended trajectory angle). Feedback information about the trajectory error is communicated by Brainsight on the bottom of the panels. S = Superior; I = Inferior; A = Anterior; P = Posterior; L = Left; R = Right.

### Electrode Implantation

#### Anchor Bolt Fixation

Once the trajectory and the entrance point were finalized, the FG arm was locked in place, allowing a craniotomy to be made along the trajectory. The pointer was first removed from the chuck on the FG arm stage and was replaced with the drill sleeve. To ensure that the drill bit does not penetrate the dura while performing the craniotomy, the distance from the top of the drill sleeve to the inner table of the skull was measured using the drill bit and ruler, in addition to the skull thickness as presented on the CT image. A stopper was placed at this distance from the tip of the drill bit and a burr hole was made using a hand drill through the drill sleeve (Fig. 15A-B). After the craniotomy, the area was irrigated with saline to make the dura visible. To secure the trajectory, an anchor bolt was seated into the burr hole and tightened into the skull using the placement wrench passed through the drill sleeve (Fig. 15C). Care was taken to not plunge down into the skull to avoid any damage to the brain parenchyma.

**Figure 15:**
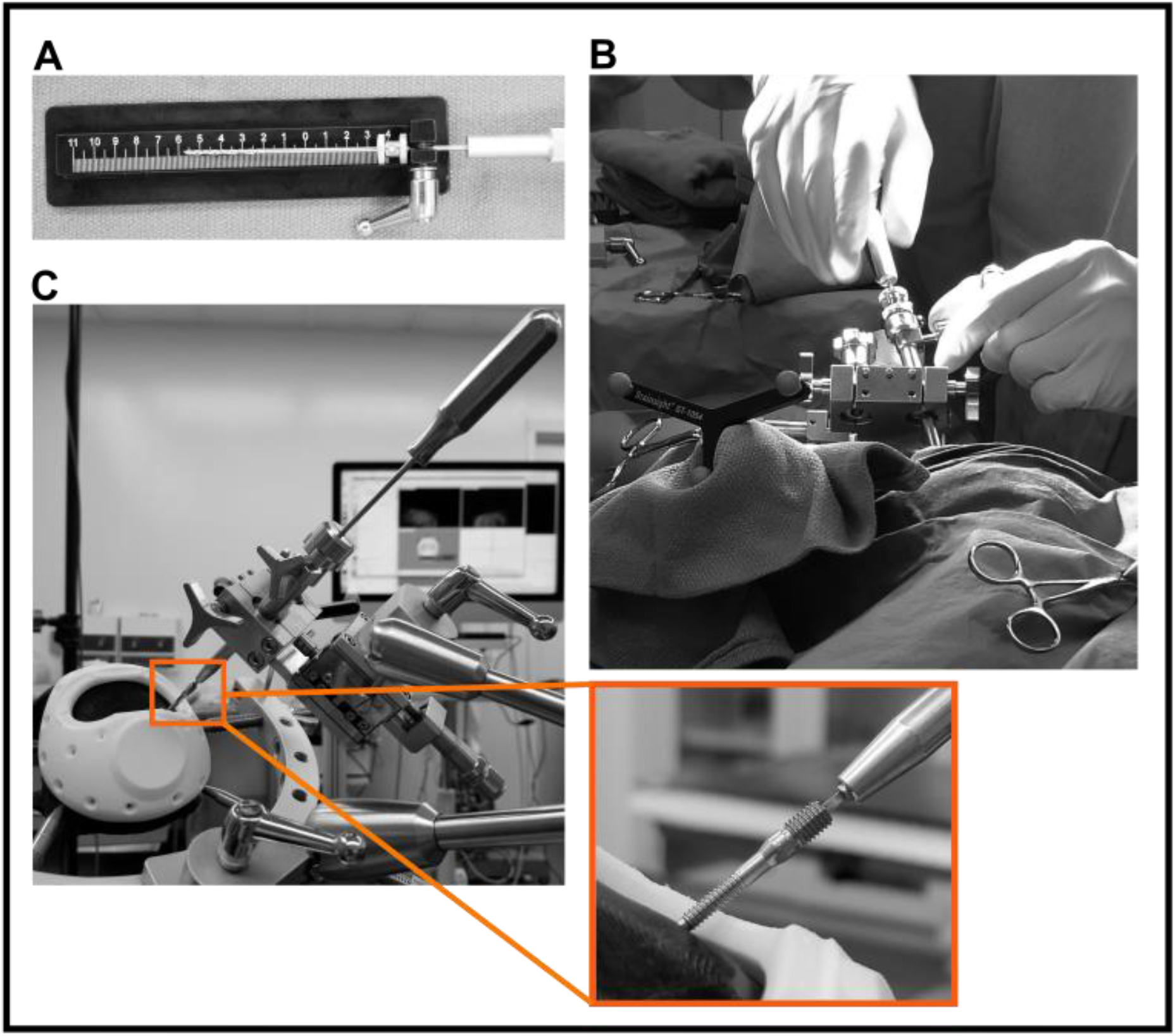
Anchor bolt insertion. **A.** Ruler is used to insert the drill bit stopper at the appropriate distance to stop at the inner table of the skull. **B.** Drill bit is fed through the drill sleeve on the stage chuck and a craniotomy is made in-line with the trajectory. **C.** After the craniotomy the drill bit is removed, and placement wrench is used to fasten the anchor bolt into the skull in-line with the trajectory. Shown here is a mock surgery with a 3D printed skull (black) and cap (white). Inset: Close up of wrench-anchor bolt coupling allowing for fastening of the bolt along the trajectory.

#### Electrode Insertion (Subject Mi)

The fixation of the anchor bolt ensured that the chosen trajectory was finalized. For the microwires to be lowered to the correct depth and placed within the structure of interest (Basolateral amygdala, BLA), the distance from the top of the anchor bolt to the target of interest was measured. The tracked pointer was placed into the chuck on the FG arm stage, positioning its tip at the top of the anchor bolt (Fig. 16A). Using the Brainsight software the distance from the tip of the pointer to the selected target was measured, and a blunt stylet was marked for insertion 5 mm above the target (Fig. 16B). This prepared a path through the brain parenchyma, allowing the guide tube to travel to above the target without deviation. Once completed, the guide tube (with a stainless-steel insert for rigidity) was lowered by hand to 5 mm above the target and fixed to the opening of the anchor bolt using dental cement (Fig. 16C). Once the cement was cured using UV-light, the insert was removed from the guide tube lumen allowing a direct path to the target. To implant the microwire brush array electrode, its microfil tube was marked to 5 mm past the guide tube length, which positioned the microwire tips within the structure of interest. The electrode was then inserted into the guide tube and lowered slowly by hand to the indicated mark and fixed to the anchor bolt and guide tube using dental cement (Fig. 16D). To obtain a ground signal, two titanium bone screws (Gray Matter Research, Bozeman, MT) were inserted into the skull, anterior to the implanted electrode (orange square, Fig. 16D). The external ground wire from each 32-channel connector was wrapped tightly around and secured to each screw (Fig. 16E). With the exception of the inserted depth of the stylet and guide tube during the second implantation, this procedure was identical in all three implantations on subject Mi. For the second procedure, the stylet and guide tube were inserted 15 mm above the target, allowing the electrode to travel through more tissue before reaching the amygdala.

**Figure 16:**
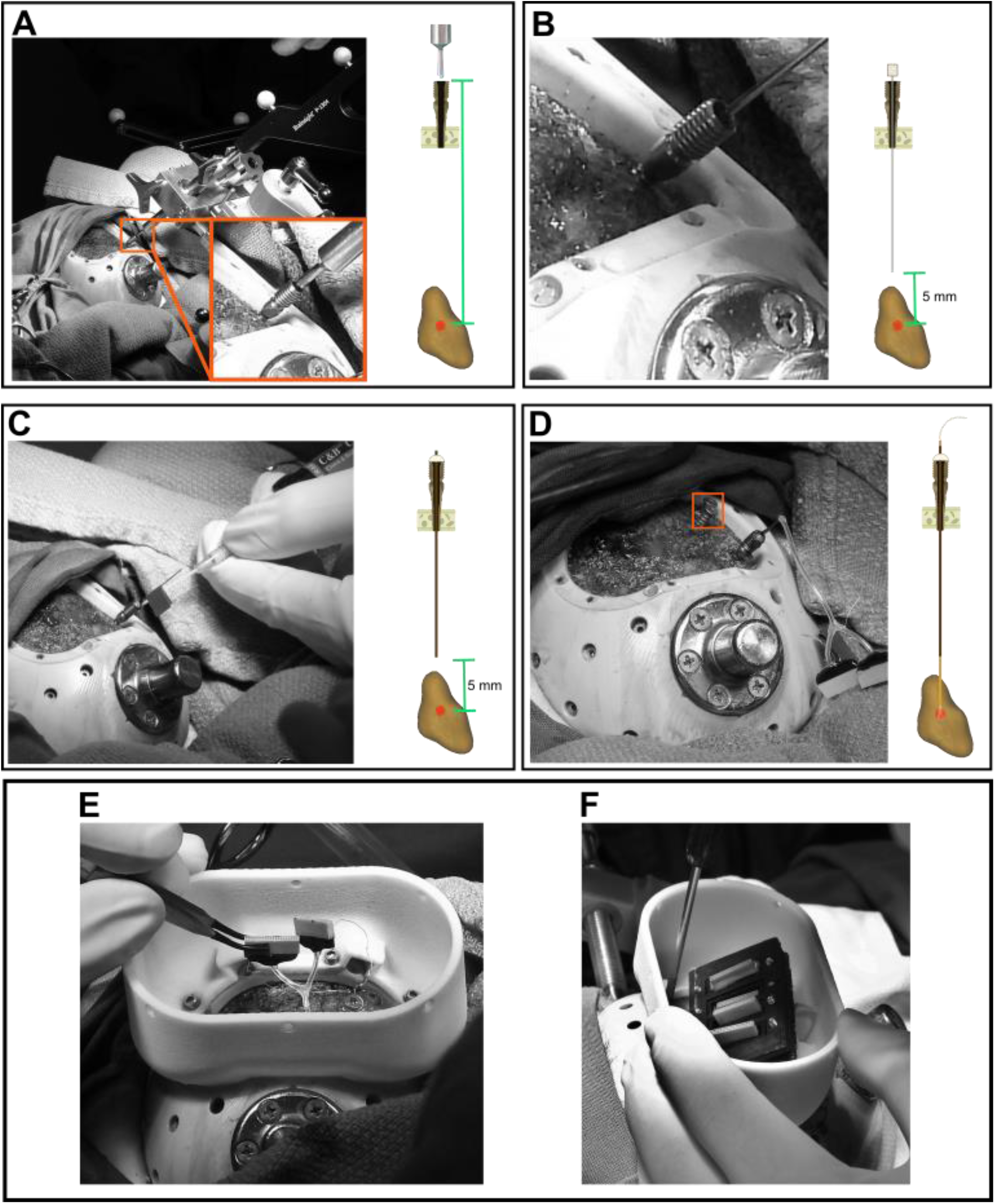
Electrode implantation and housing. **A.** Distance from top of the anchor bolt to target of interest was determined using the pointer tip in Brainsight. Inset: Close-up of pointer tip secured above the anchor bolt opening. Schematic: Pointer placed on top of the anchor bolt (cross-section) secured into the skull and distance to target (red circle) in the BLA (amygdala reconstruction in brown) is shown by the green marker. This distance is variable depending on the selected trajectory. **B.** Blunt stylet passed through the anchor bolt to prepare the trajectory for guide tube insertion. Schematic: Stylet was stopped 5 mm above the target. **C.** Guide tube was inserted into the anchor bolt and fixed using dental cement. Stainless steel insert can be seen marked with the “flag”. Schematic: Guide tube inserted to 5 mm above the target and fixed with dental cement (white). **D.** Electrode is lowered by hand to the target and fixed to the anchor bolt-guide tube assembly with dental cement. Orange square indicates grounding screws fastened into the skull Schematic: Electrode microwires positioned in the structure of interest after passing through the guide tube. **E.** Ground wires were secured to their respective screws and connectors were secured in their housing (**F**).

To ensure safe housing for the electrode and the connectors, the bowl enclosure was secured onto the cap using screws, covering the implanted region. Skull surface was covered with silicone (Kwik-Sil; WPI Surgical Instruments, Sarasota, FL), covering the ground wires while ensuring some slack to avoid snapping of the wires. Lastly, each 32-channel connector was cemented into a single connector holder and screwed onto the bridge housing, which held up to four connectors (Fig. 16E-F). This bridge housing was then cemented onto the bowl enclosure wall. This housing allowed easy access for connecting the headstage during each recording session while providing a secured place for the connectors between sessions.

#### Electrode Insertion Using Microdrive (Subject Ma)

The anchor bolt was secured in-line of the trajectory using the placement wrench. Dental cement was applied to the base of the anchor bolt to further secure it to the skull (Fig. 17A). A stainless-steel grounding screw was inserted subsequently after (Fig. 17B). After determining the distance from the top of the anchor bolt to the target, the microdrive was installed. The microdrive was screwed onto the anchor bolt threads and dental cement was subsequently applied to the underside junction with the anchor bolt (Fig. 17C-D). The microdrive clamp was left in an open position to allow access to the anchor bolt opening. A blunt stylet was marked and inserted to 2 mm above the target (Fig. 18A), followed by insertion of a guide tube with a stainless-steel insert (Fig. 18B). Once the guide tube reached 2 mm above the target, it was secured to the anchor bolt using dental cement and the insert was removed, allowing the guide tube to be cut level with the anchor bolt opening (Fig. 18C). The electrode was then marked and lowered through the guide tube by hand to 1 mm above the guide tube’s distal opening (3 mm above the target; Fig. 18D). The electrode clamp was then closed, holding the electrode in place. The electrode was then slowly lowered 3 mm using the microdrive, positioning the microwires within the target (Fig. 18E). The grounding wire was tightly wrapped onto the grounding screw and covered with dental cement (Fig. 19A). The microdrive cover was placed over the body, in a top-down manner, using the outlet slit to accommodate the connector and grounding wires, and screwed onto the drive body (Fig. 19B). Finally, the connectors were cemented into the connector holders, which were screwed onto the connector bridge. The bowl enclosure was screwed on, and the bridge was cemented onto its walls through the dental cement pocket (Fig. 19C). The enclosure lid was screwed onto the bowl, completing the implantation procedure.

**Figure 17:**
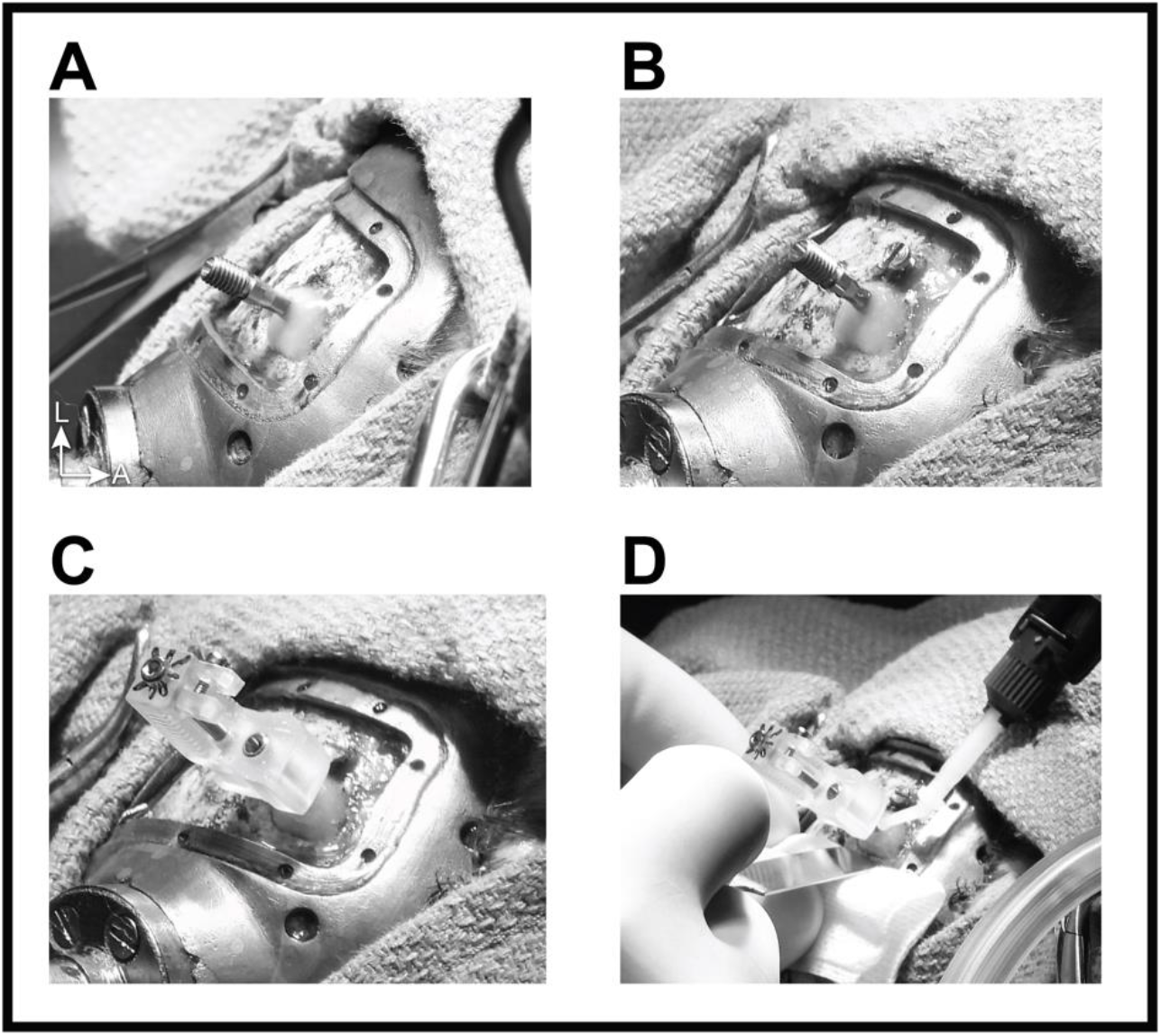
**A.** Securing the anchor bolt to the skull using dental cement. **B.** Grounding screw insertion into the skull. **C.** Screwing on the assembled microdrive onto the anchor bolt with the electrode clamp in open position. **D.** Reinforcing the microdrive hold onto the anchor bolt using dental cement at the junction underneath. S = Superior; I = Inferior; A = Anterior; P = Posterior; L = Left; R = Right.

**Figure 18:**
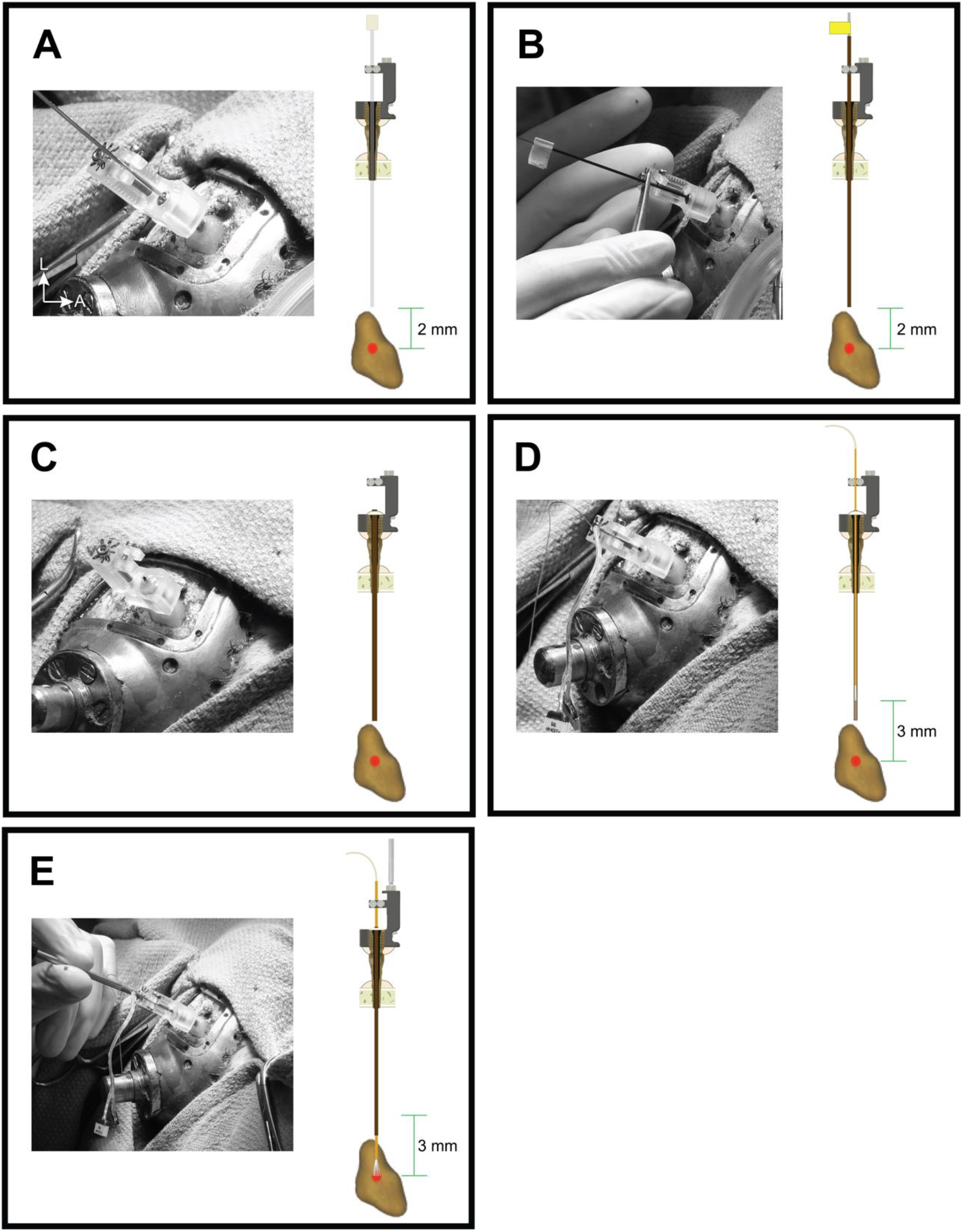
**A.** With the microdrive clamp open, the stylet was lowered by hand through the anchor bolt to 2 mm above the target. **B.** The guide tube, with an insert (yellow tape used as the stopper), was lowered by hand to 2 mm above the target. **C.** Guide tube cemented to the anchor bolt and cut leveled with the anchor bolt opening. **D.** The electrode was lowered to 3 mm above the target and secured by the clamp. **E.** Electrode lowered 3 mm with the microdrive to reach the target. Note: Schematics are not to scale and are for demonstration purposes only. S = Superior; I = Inferior; A = Anterior; P = Posterior; L = Left; R = Right.

**Figure 19:**
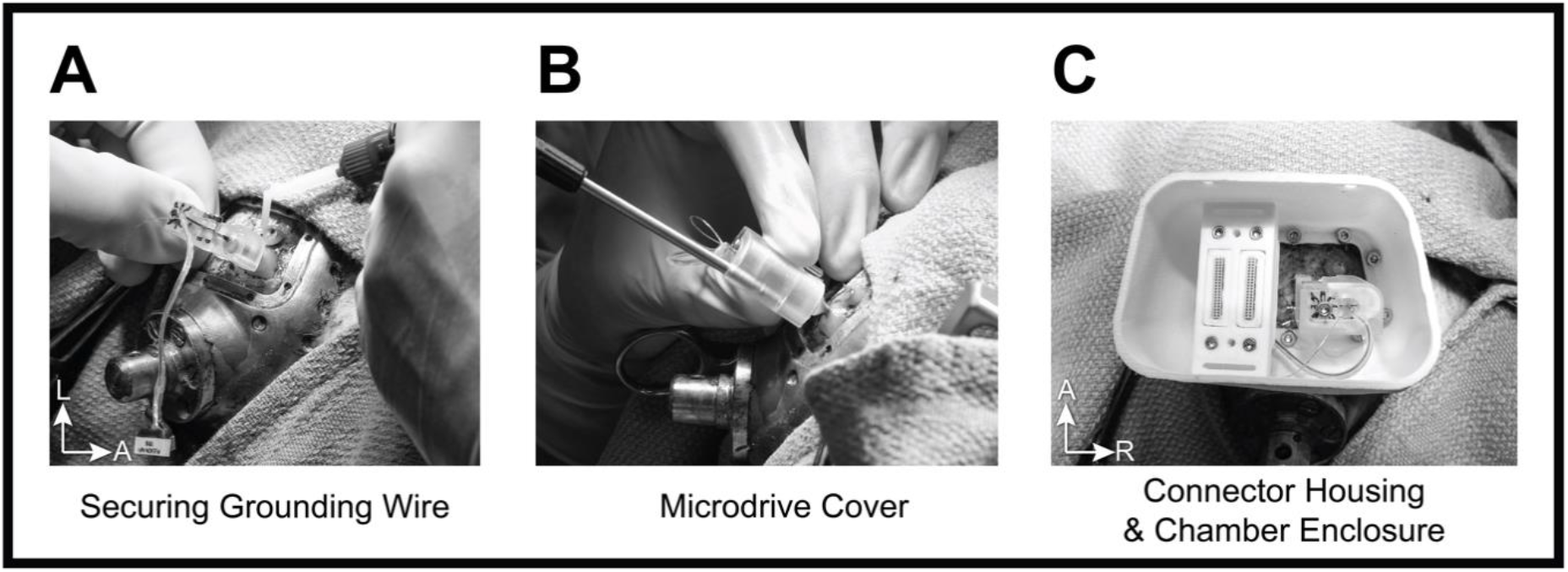
**A.** The ground wire was wrapped tightly around the grounding screw and covered using dental cement. **B.** A screw was used to secure the microdrive cover to the body. **C.** Final bowl enclosure assembly. The bowl enclosure was secured to the cap with screws. The connectors were secured within their housing and fixed to the walls of the bowl enclosure by applying dental cement through the horizontal slits. The lid (not shown) subsequently covers the assembly. S = Superior; I = Inferior; A = Anterior; P = Posterior; L = Left; R = Right.

## Results

### Targeting error

The NHPs were monitored daily post-implantation, and no complications were noted based on behavioural observations. Two weeks post-implantation, a CT scan (154 µm resolution) of the subjects were obtained to determine the position of the recording electrode (Fig. 20A i-ii, Fig. 21A-B i, Fig. 22A i-ii, Supplementary Figure 3A i-iii). To assess the targeting error, the post-electrode-implantation CT was co-registered with the pre-cap CT, which as previously described was aligned with the T1 MRI sequence. The protocol for this manual rigid point-based co-registration was similar to that of pre-to-post cap CT co-registration previously described using 3D Slicer (Fedorov et al., 2012). The deviation of the implanted electrode from the intended target was assessed in all three anatomical axes relative to the orientation of electrode, and a single metric of Euclidean distance from the target (total error) was reported (Table 1, Fig. 20B-C, Fig. 21A & C, Fig. 22B-C, Supplementary Figure 3B-C).

**Table 1:** Targeting errors for all 4 implantations performed (across subject Mi and Ma): Implant number refers to the specific electrode implantation performed on each subject. Errors for each axis are represented separately. Total error (vector of error) indicated is the Euclidean distance of the electrode tip in the post-implant CT to the desired target chosen pre-implantation. This is calculated for each implant and the average error separately using the axes outlined. Note*: Implantation 2 on subject Mi is not used in the average error calculation due to the electrode deviation during implantation. Med./lat.: Medial-Lateral axis, Ant./post.: Anterior-Posterior axis, Sup./inf.: Superior-Inferior axis.

| **Subject** | **Implant number** | **Med./lat.(x)** | **Ant./post.(y)** | **Sup./inf.(z)** | **Total error (Euclidean distance)** |
| --- | --- | --- | --- | --- | --- |
| Mi | 1 | 0.5 mm | 0.8 mm | 0.8 mm | 1.2 mm |
|  | 2* | 0.5 mm | 1.6 mm | 5.0 mm | 5.3 mm |
|  | 3 | 0.5 mm | 0.9 mm | 1.0 mm | 1.4 mm |
| Ma | 1 | 2.2 mm | 0.7 mm | 0.3 mm | 2.3 mm |
| Average error* |  | 1.1 mm | 0.8 mm | 0.7 mm | 1.6 mm |

**Figure 20:**
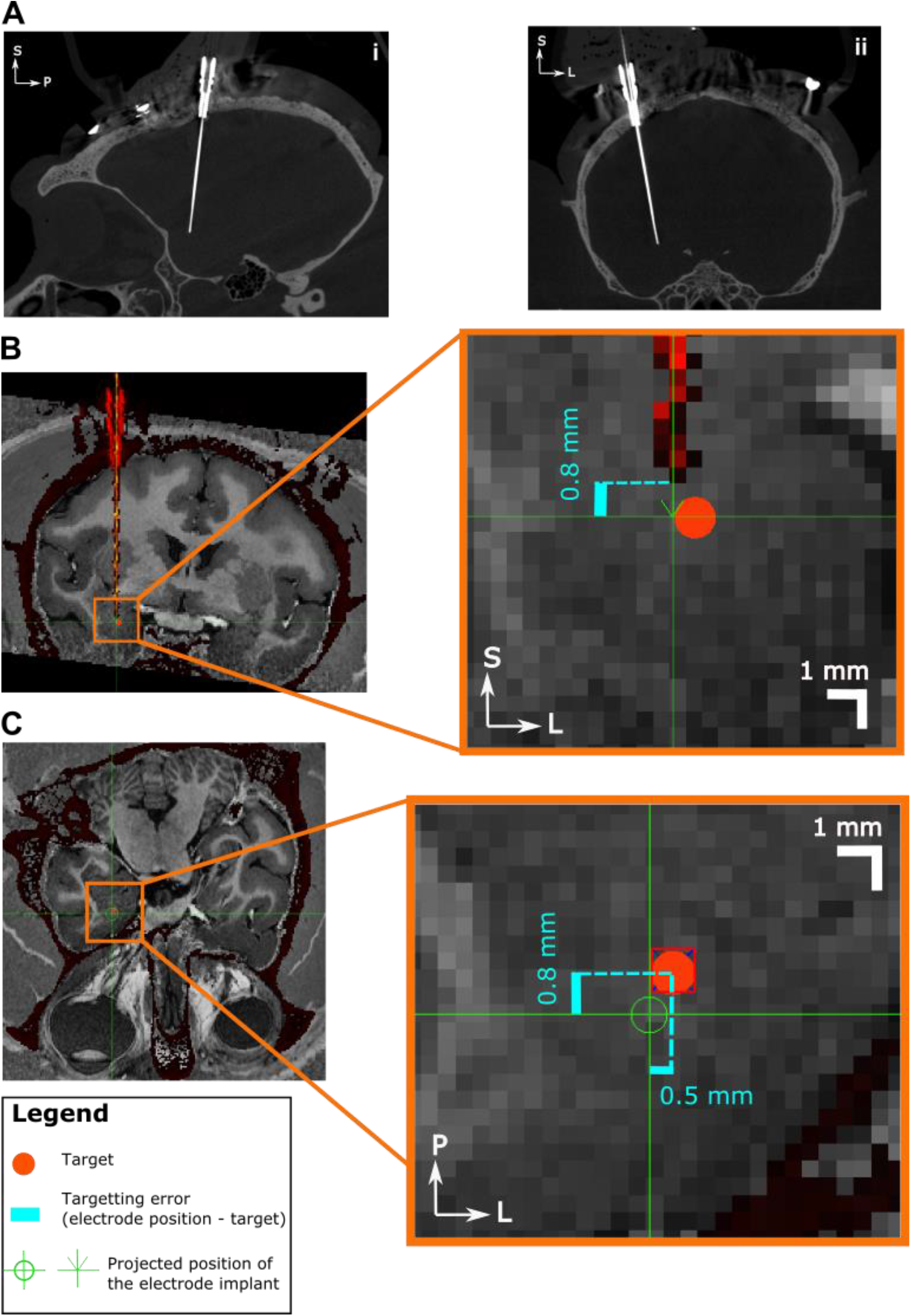
Targeting error for the first implantation for subject Mi. **A.** Sagittal (i) and coronal (ii) views respectively, of post-implantation CT scan. B) Coronal view of the intensity thresholded image of the CT, co-registered with subject MRI for assessment of targeting error in the superior-inferior axis. Implanted electrode and anchor bolt are pictured red. **C.** Transverse view of the implantation, with the electrode projected tip to enable targeting error measurement in the two axes (anterior-posterior and medial-lateral axes). Skull is shown as dark red. Insets: Magnified segments of MRI images to locate electrode tip and quantify targeting error. Actual electrode shown intensity thresholded in red pixels. Projected electrode position is shown in green. Figure legend describes inset graphic details. S = Superior; I = Inferior; A = Anterior; P = Posterior; L = Left; R = Right.

**Figure 21:**
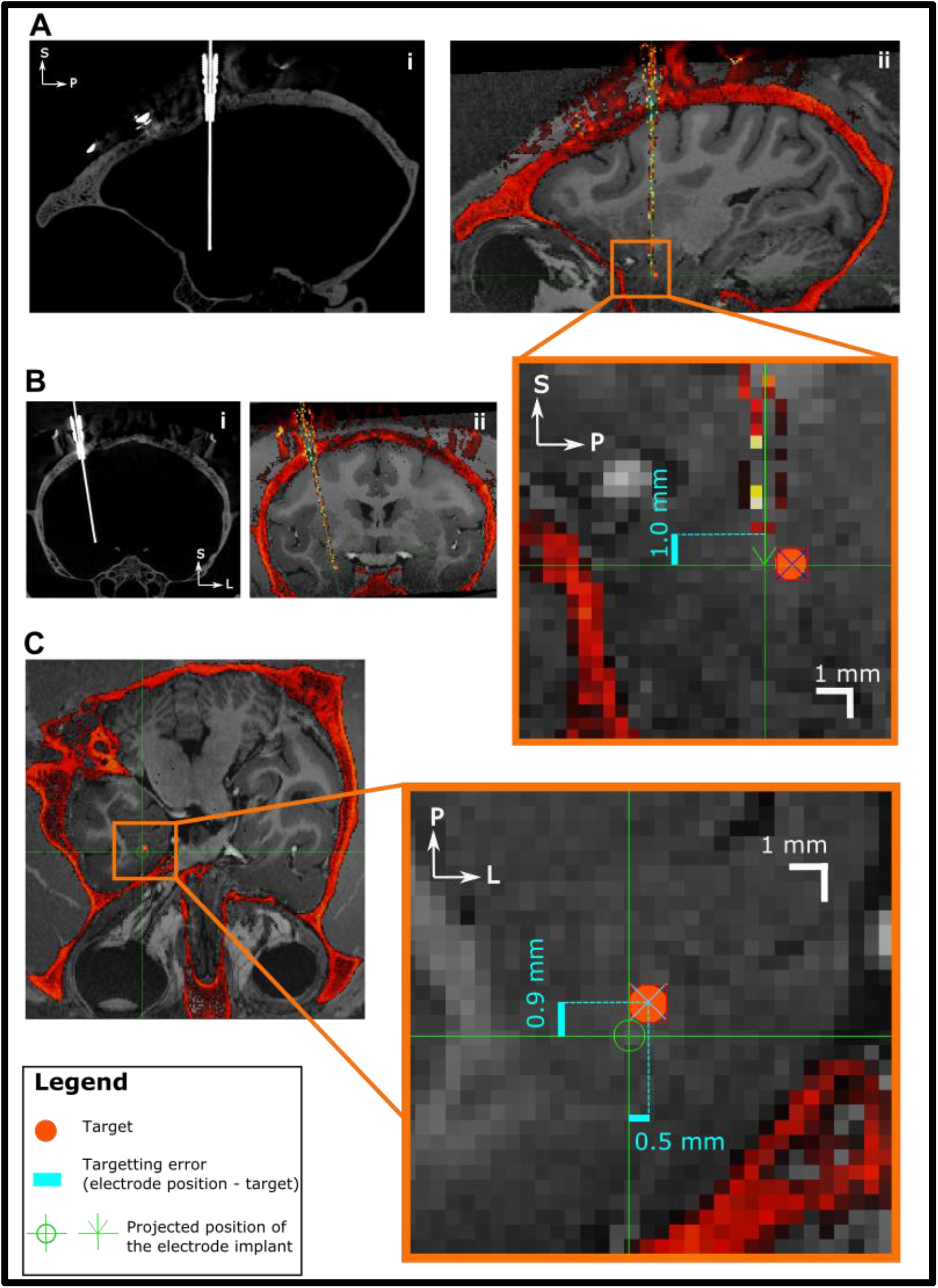
Targeting error for the third implantation for subject Mi. **A**-**B.** Sagittal and coronal views respectively, of post-implantation CT scan (i) and thresholding of the CT co-registered with subject MRI (ii) for assessment of targeting error. **C.** Transverse view of the implantation, with the electrode projected tip to enable targeting error measurement in the two axes. Skull is shown in red in all intensity thresholded images. Insets: Magnified segments of MRI images to locate electrode tip and quantify targeting error. Actual electrode shown intensity thresholded in red pixels. Projected electrode position is shown in green. Figure legend describes inset graphic details. S = Superior; I = Inferior; A = Anterior; P = Posterior; L = Left; R = Right.

**Figure 22:**
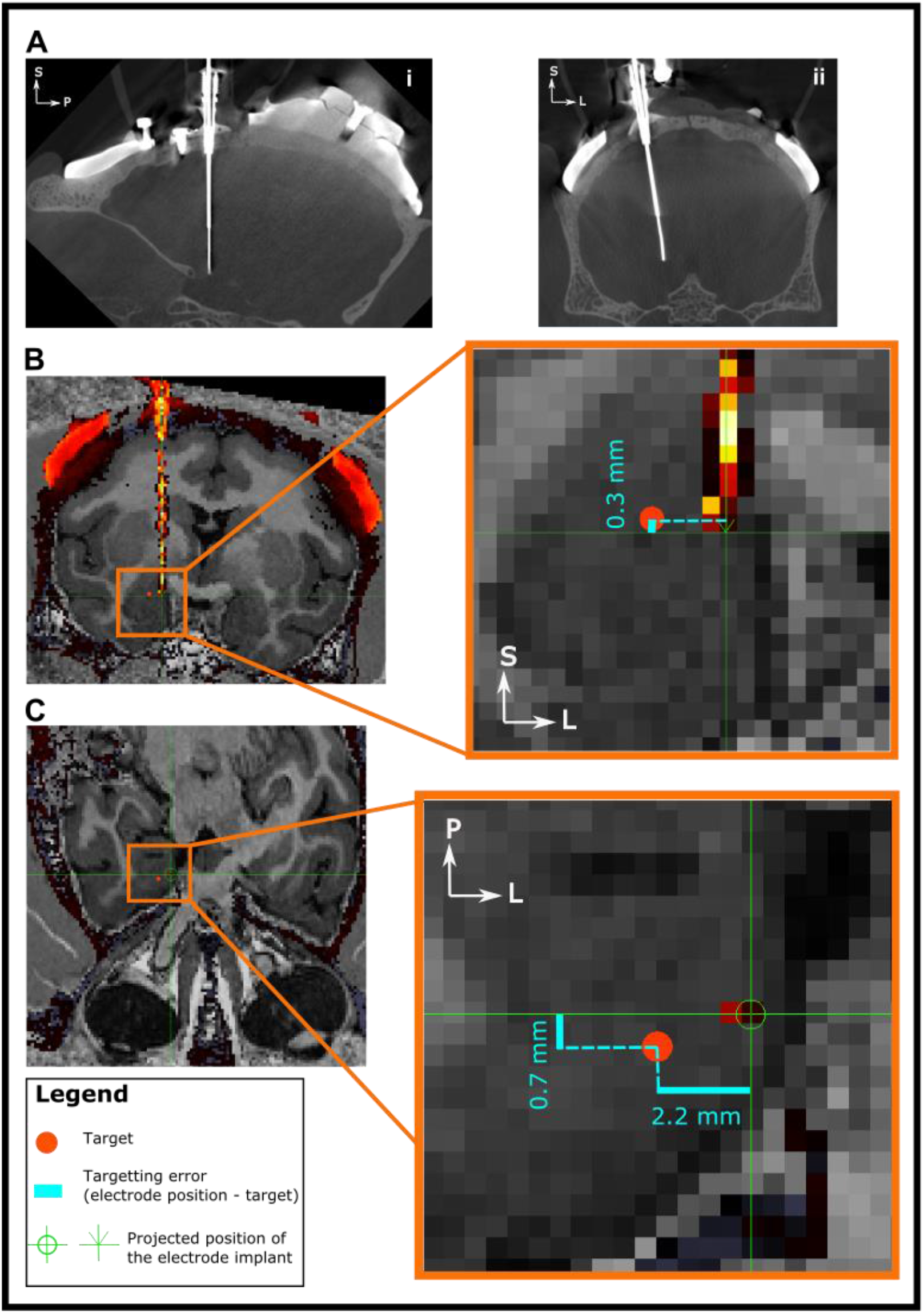
Figure 16: Targeting Error for monkey Ma. **A.** Sagittal (i) and coronal (ii) views of the post-implantation CT scan in-line with the electrode. **B-C.** Coronal and transverse view of the post-implantation CT respectively, co-registered with subject MRI. CT is intensity thresholded (red) to enable visualization of electrode position to assess targeting error. Magnified insets: Targeting error quantification of the implanted electrode (intensity thresholded red pixels) is shown in superior-inferior (B inset), medial-lateral (C), anterior-posterior (C) axes. Figure legend describes inset graphic details. S = Superior; I = Inferior; A = Anterior; P = Posterior; L = Left; R = Right.

A total of three implantations were performed on monkey Mi. Unlike the procedures described in Figure 16 for our first and third implantation, the guide tube during the second procedure was placed 15 mm above the target, and the electrode was lowered to the target. This was performed to increase the splay of the microwires as they travel through more brain tissue. Unexpectedly, the microwire tips bent before reaching our basolateral amygdala target, as can be seen in post-implantation CT images of supplementary Figure 3. This resulted in a total error of 5.2 mm, with the errors resulting from deviations in the following anatomical axes of the trajectory: 0.5 mm in the medial-lateral axis, 1.6 mm in the anterior-posterior axis, and 5 mm in the superior-inferior axis. As this unusual deviation of the microwires from the intended trajectory was not a result of our targeting methodology, here we will omit this implantation from our calculations of total targeting error. We have presented this case as a caution for future investigators looking to replicate this method. Although factors resulting in this deviation cannot be confirmed, we hypothesize that one possibility is the microwires attempting to pierce the pia matter after entering a subarachnoid cistern in the trajectory path. However, the probability of this is low given that these anatomical spaces were not visible in the trajectory path on the MRI. A second possibility is the bending of the microwires as a result of change in the tissue resistance, when transitioning from grey to white matter. If any differences exist in the density of these structures, it could cause the deflection of the microwires as we have observed.

The total error for the first implantation in monkey Mi (Figure 20) was 1.2 mm, with the errors distributed in the following anatomical axes of the trajectory: 0.5 mm in the medial-lateral axis, 0.8 mm in the anterior-posterior axis, 0.8 mm in the superior-inferior axis. The third implantation (Figure 21) resulted in a total error of 1.4 mm was measured in the three axes as follows: 0.5 mm in the medial-lateral axis, 0.9 mm in the anterior-posterior axis, and 1.0 mm in the superior-inferior axis. Due to the resolution of the CT image, determining the exact position of individual microwires is not possible. Therefore, these errors represent the best estimate of the microwire positions, assuming a uniform splay of all 64 microwires. For monkey Ma, a total error of 2.3 mm was observed from the electrode tip to the target. The error was distributed in the three anatomical axes of the trajectory as follows: 2.2 mm in the medial-lateral axis, 0.7 mm in the anterior-posterior axis, and 0.3 mm in the superior-inferior axis. In summary, the average total targeting error of our three subcortical implantations across two animals was 1.6 mm.

### Neural recordings

One-week post-implantation, the collection of neural recordings began as the animals performed various behavioural tasks seated in front of a monitor. Cerebus Neural Signal Processor (Blackrock Microsystems) was used for neural data collection, digitized at 16 bit and sampled at 30kHz. The spike waveforms were detected online by thresholding at 2.5 to 3.5 multiplier of the root mean squared signal energy and the units were manually sorted offline using Plexon Offline Sorter (Plexon Inc.). Figure 23 shows waveforms and their associated PCA cluster of five single unit examples sorted from implantation two and three in monkey Mi during the four months of recording. The ground connection for implantation one in monkey Mi was damaged during post-implantation recordings due to housing issue, leading to faulty recordings, and therefore are not included here. Recordings in subject Ma did not yield any single or multi-unit recordings and are therefore not shown here. Monkey Mi’s electrode implants were recorded from during a 7-month period without complications before the termination point was reached. Monkey Ma’s electrode implant lasted a total of 13 months before health complications resulted in euthanasia of the animal.

**Figure 23:**
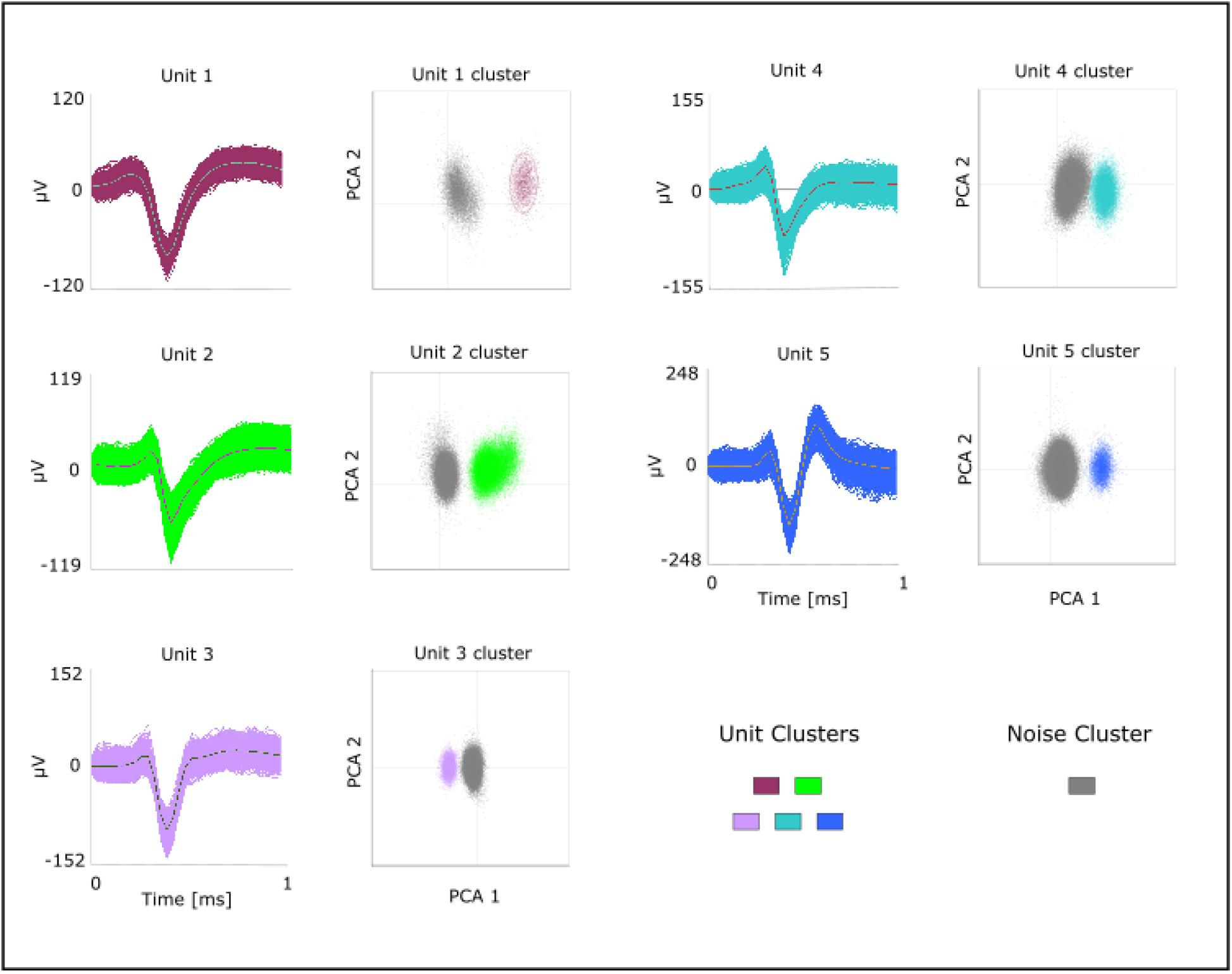
Single-unit activity from subject Mi (implantation two and three). **A.** Shown are 6 examples of single activity recorded during the 3-month recording period. Each color denotes a different unit, and the black line shows the average wave form of each unit. **B.** Clustering of each unit and noise using 2 PCA features of the waveforms. Color corresponds to units in A.

## Discussion

In this paper we have described our methodology for chronic and semi-chronic implantation of microelectrode arrays in the amygdalae of two macaque monkeys using a neuronavigation procedure. The average targeting error of 1.6 mm achieved for our three implantations is within the range reported in previous literature targeting deep brain structures. This error is lower than or equal to what has been previously reported in human neuronavigation procedures (Bjartmarz & Rehncrona, 2007; Bradac et al., 2017; Holloway et al., 2005; Sharma et al., 2014). Our error is also lower in comparison to targeting accuracies reported in neuronavigation brain biopsy surgeries in other animal models (e.g. dogs) (Chen et al., 2012; Long et al., 2014). Although in NHPs, higher targeting accuracies using neuronavigation for injections of deep regions, such as the cerebellar dentate nucleus, have been reported (Wetzel et al., 2019), the desired target region used for assessment were also larger than our 1 mm diameter target in the basolateral nucleus of amygdala. Lastly, our targeting accuracies are lower in comparison to the improved MRI-guided stereotactic procedures targeting deep regions (Walbridge et al., 2006). However, the targeting accuracies in these procedures were determined by comparison of pre-to post-operative MRI, with slice thickness of 1-3mm, in comparison to our use of CT imaging (154-micron resolution). This discrepancy in accuracy between frame-based (stereotaxic) and frameless (neuronavigation) systems is not often reported in deep brain procedures in humans (Dhawan et al., 2019; Holloway et al., 2005). It is worth noting that comparison between targeting errors across procedures can be difficult for several reasons. Despite using the same implantation technique (e.g. neuronavigation), the implantation operations can have different workflows leading to targeting inaccuracies arising from different factors (Li et al., 2016). The declared target of interest, to which trajectory deviations are compared, also tend to vary in different studies ranging from a cerebellar nuclei (Wetzel et al., 2019) to specific coordinates within a deep brain structure (Chen et al., 2012). Lastly, measurements of the targeting errors are performed using various methods ranging from histological analysis to post-implantation MRI with varying slice thickness, or CT imaging which can introduce a measurement error during assessment of targeting due to MRI-CT image fusion errors (Roth et al., 2018).

To improve the targeting accuracy of our implantations, we sought to make use of cranial cap screws as fiducial markers for image-subject registration. Two of the major contributors to targeting error in neuronavigation implantations are errors in locating fiducial markers, and their physical displacement (e.g. movement of skin adhesive fiducials) from the time of subject imaging to image-subject registration during surgery (Wang & Song, 2011). As cranial cap screws are stable and easily accessible landmarks, they allowed for reduction of such errors in the registration process. Future improvements for this fiducial system could include a method for reduction of the fiducial localization error (FLE) by restricting the tracked pointer’s degrees of freedom of movement when localizing a fiducial. This would ensure that the tracked pointer remains in-line with the indicated fiducial during the point-to-point registration process. To further reduce FLE, multiple samples of each fiducial point could be collected, allowing for the use of an average of samples for localization of each fiducial point during registration. This would reduce the inherent variability in manual localization of fiducials (Woerdeman et al., 2007) as previously reported. An additional benefit of using cranial cap screws as fiducial markers is their distribution across the subject’s skull. Improper distribution or configuration of fiducial makers relative to the surgical field (i.e. target position) can further contribute to registration errors which will increase the targeting error (Wang & Song, 2011; West et al., 2001). Previous studies using dental imprint platforms for mounting of fiducial markers do not allow for their placement close to target position (Johnston et al., 2016), while other approaches using skull mounted fiducials have a limited distribution (Frey et al., 2004). Our cranial cap screw distribution provides a fiducial set, of which a subset can be chosen for registration depending on the target position of interest. Furthermore, it may be possible to reduce the number of selected fiducials without compromising the registration accuracy, resulting in reduced surgical time. Quantitative assessment of the effect of the point-to-point registration transformation on displacement of the true target position (i.e. target registration error, TRE) will aid in diagnosing of future targeting errors (Fitzpatrick, 2009; Fitzpatrick et al., 1998). Lastly, although placement of the electrodes by hand was performed to replicate microwire implantation protocols in humans (Misra et al., 2014), it likely contributed to our targeting errors in the superior-inferior axis (Table 1) of the trajectory due to lack of control over the depth of insertion. Alternatively, the use of a micromanipulator during implantation would provide greater control and precision, potentially improving the targeting performance and producing minimal tissue damage.

In addition to the presented targeting approach, we also demonstrated a technique for implantation of a minimally invasive chamber for chronic and semi-chronic recordings. We were able to collect chronic single unit recordings in subject Mi for duration of four months, albeit with low yield. Recordings in subject Ma did not yield single or multi-units. Our choice of electrode for this procedure was informed by previous studies using microwire bundles for long-term chronic recordings with adequate yield (Bondar et al., 2009; McMahon et al., 2014; Porada et al., 2000). However, we were unable to replicate these results with our implantation protocol. One mode of failure in microwire recordings is suggested to be a lack of splay or outspreading of individual microwires in the bundle during implantation to allow for successful unit recordings (Babb et al., 1973; Misra et al., 2014). Due to the size of the individual microwires (12 μm in diameter), we were unable to verify this lack of microwire splay using post-implantation CT, unlike what has been previously demonstrated in human implantations (Misra et al., 2014) using larger microwires. However, we did observe a high cross-correlation of signals across the channels, suggestive of microwires remaining in a bundle in the neural tissue. Despite the low yield, the implantation protocol described here can be implemented using any single shank recording electrode. Future work using this protocol could utilize the current multi-electrode laminar probes available for deep brain recordings in NHPs (Pomberger & Hage, 2019).

The use of the small anchor bolt chamber proved to be advantageous as only a small burr hole was required, reducing the risk of infection and other potential complications. This method is easily implementable by any research group as the components used in the protocol are readily commercially available. Although not demonstrated in this protocol, the small outer diameter of this chamber allows for implantation of several adjacent trajectories with differing implantation angles, targeting different structures. The use of a mounted microdrive could add further longevity and flexibility to such implantations. In our experience in designing such microdrives, we suggest the use of a clamping mechanism for attachment of the microdrive to the anchor bolt to ensure greater stability. Alternatively, addition of a scaffolding to the design of the microdrive-anchor bolt junction could allow for imbedding of this junction with dental cement for further adhesion.

In summary, we have described a targeting and implantation methodology for conducting deep brain chronic and semi-chronic electrode array recordings in the macaque monkey. Using a combination of brain navigation techniques, available tools from human and animal neurosurgery, and 3D printing technology, we have a provided a protocol that allows us to accurately target amygdala nuclei through a minimally invasive and commercially available recording chamber. As accurate localization of specific brain structures is becoming increasingly important in systems neuroscience, technologies for accurate targeting of deep brain regions are rapidly evolving. Robot-assisted neuronavigation systems is one such technology that is becoming increasingly popular and allows for reduction of targeting error beyond the gold standard of image-guided stereotaxy (Ose et al., 2022). Additionally, this technology allows for a reduction in the surgical time and provides better patient outcome as presented in the human neurosurgery literature (Fomenko & Serletis, 2018; Joswig et al., 2020). The greater accuracy and precision in targeting structures will allow for a finer study of brain circuits involved in cognitive and behavioural processes in NHPs, shedding a light on their potential role in pathophysiological conditions of human disorders.

## Appendix A: Microdrive CAD Software and Printing Information

A Form 2 SLA printer by Formlabs company was used. The Clear V4 Resin (FLGPCL04) material by Formlabs was used for printing and layer thickness of 0.025mm was chosen to optimize the resolution. When preparing material for print, the designs were placed upright in the software and a raft and support was autogenerated for each object. This allowed the touchpoints of the supports to be generated at locations of the drive body that once taken off, would not obstruct the movement of the drive clamp which was fitted in the body. After printing, parts were washed in 90% Isopropyl Alcohol for 30-60 minutes in Formlabs Wash Station. Although curing of the printed pieces is recommended for this resin, we found that any post processing (drilling or tapping) was best done before the curing as the material was more malleable.

## Acknowledgements

We thank Kim Thomaes, Kristy Gibbs, and Rhonda Kersten for their help with animal care and surgical procedures in their role as veterinarian technicians. Joseph Umoh and Trevor Szekeres for CT and MRI imaging of the animals respectively. Mohamad Abbass for his thoughts on possible causes of implantations deviation. Diego Fernando Buitrago Piza for capturing of surgical images. Andrew Pruszynski and Susheel Vijayraghavan for their help in troubleshooting the recordings. Stephen Frey from Rogue Research Inc. for his continuous support in using the Brainsight neuronavigation system. Nicolas Alba and the team at Microprobes for troubleshooting of the Microwire Brush Array (MBA) implantations. Kenji Koyano and David Leopold for their correspondence and help in troubleshooting of MBA implantation techniques and yield. This work was supported by Canadian Institute of Health Research Project Grant (CIHR); Natural Sciences and Engineering Research Council of Canada (NSERC); Canada Foundation for Innovation (CFI); Ontario Graduate Scholarship (OGS).

## Supplementary Material

**Supplemental Figure 1:**
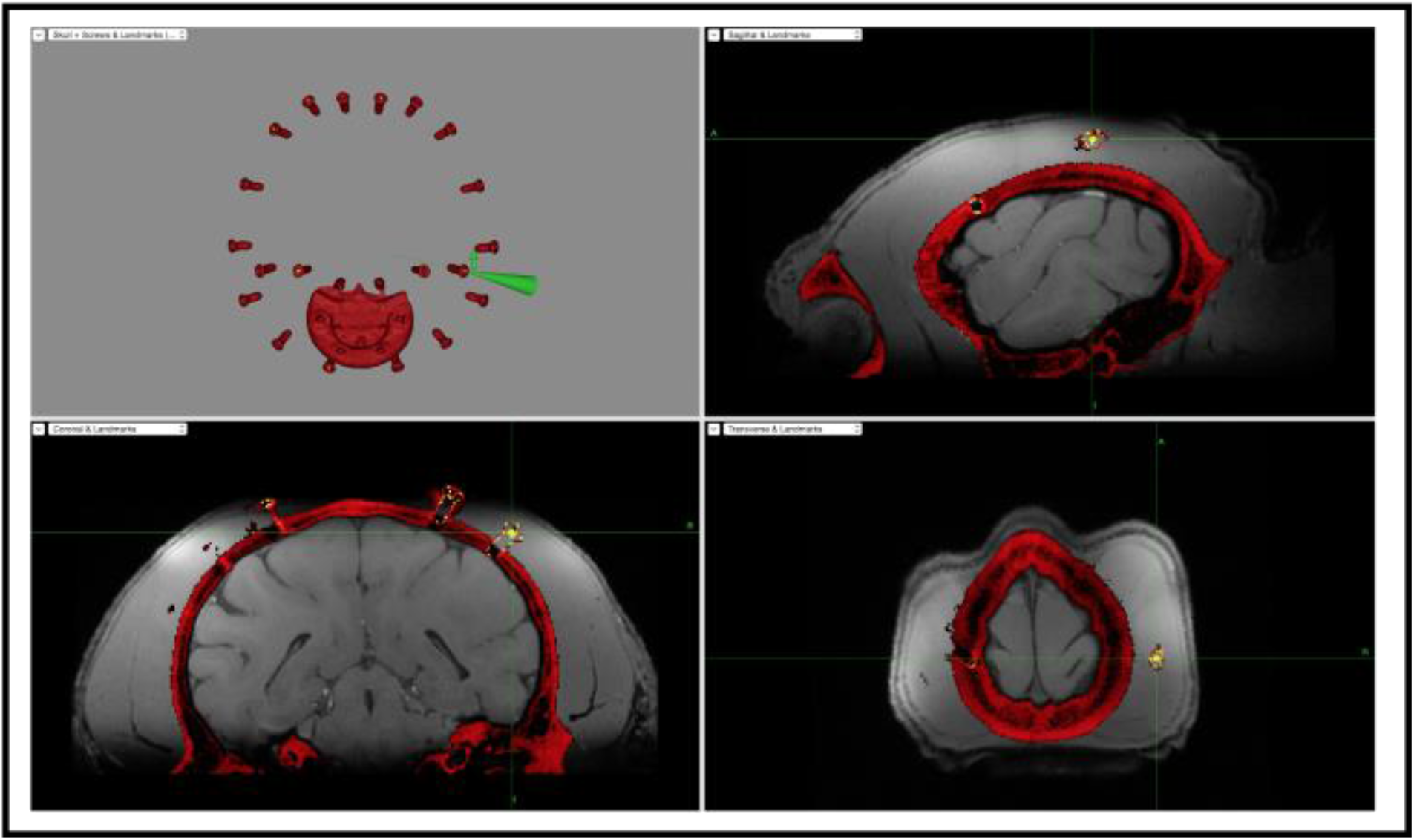
Validation of subject-image registration. View of the Brainsight software screen as the pointer is placed at various locations on the cap after registration is completed. This is performed to assess the quality of the registration. The skull in the CT scan is intensity thresholded in red.

**Supplemental Figure 2:**
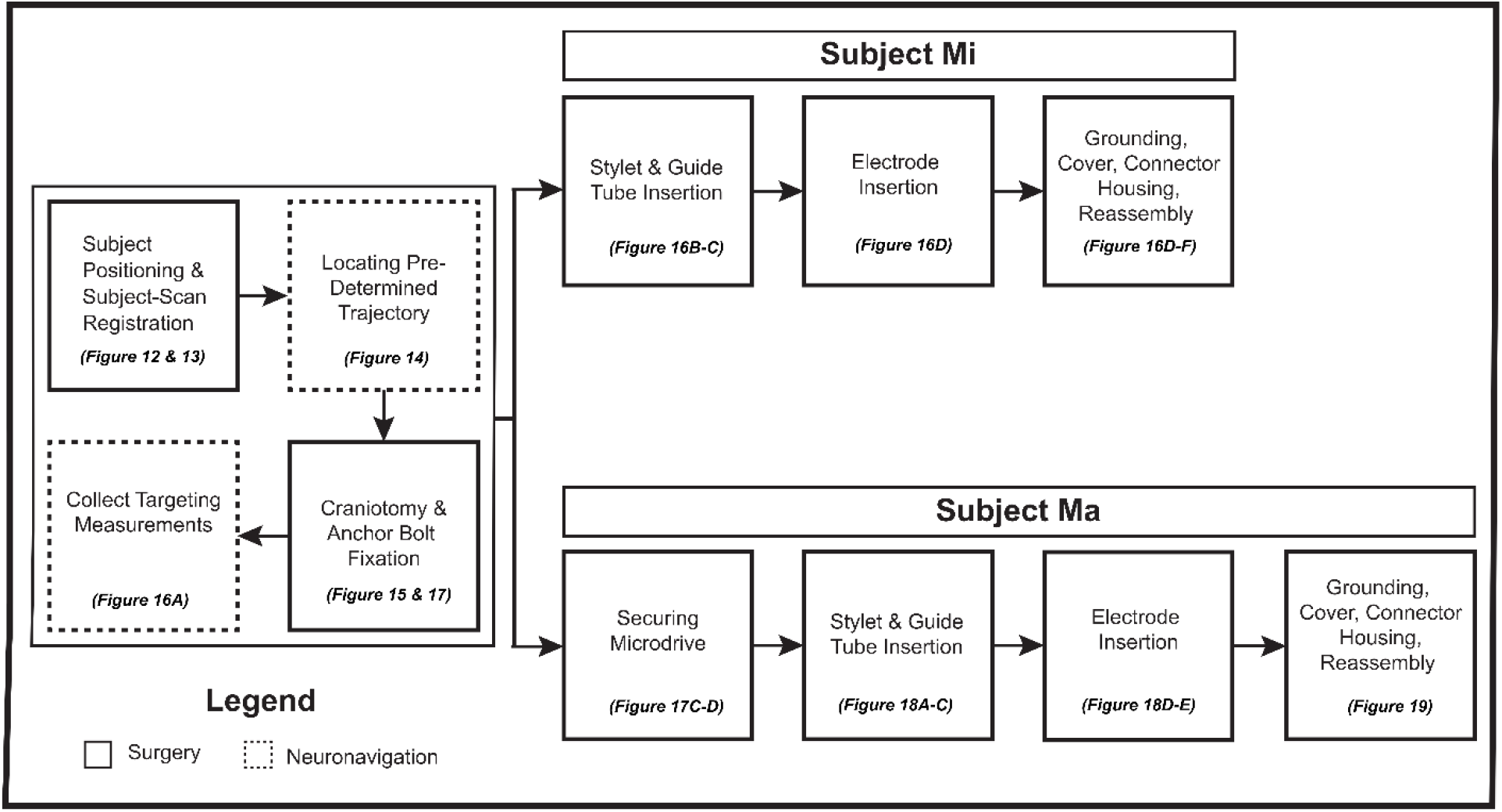
Overview of surgical protocol. The left box presents methods common to both procedures. Subject Mi received a chronic implantation, while subject Ma received the semi-chronic implantation with the use of a microdrive.

**Supplementary Figure 3:**
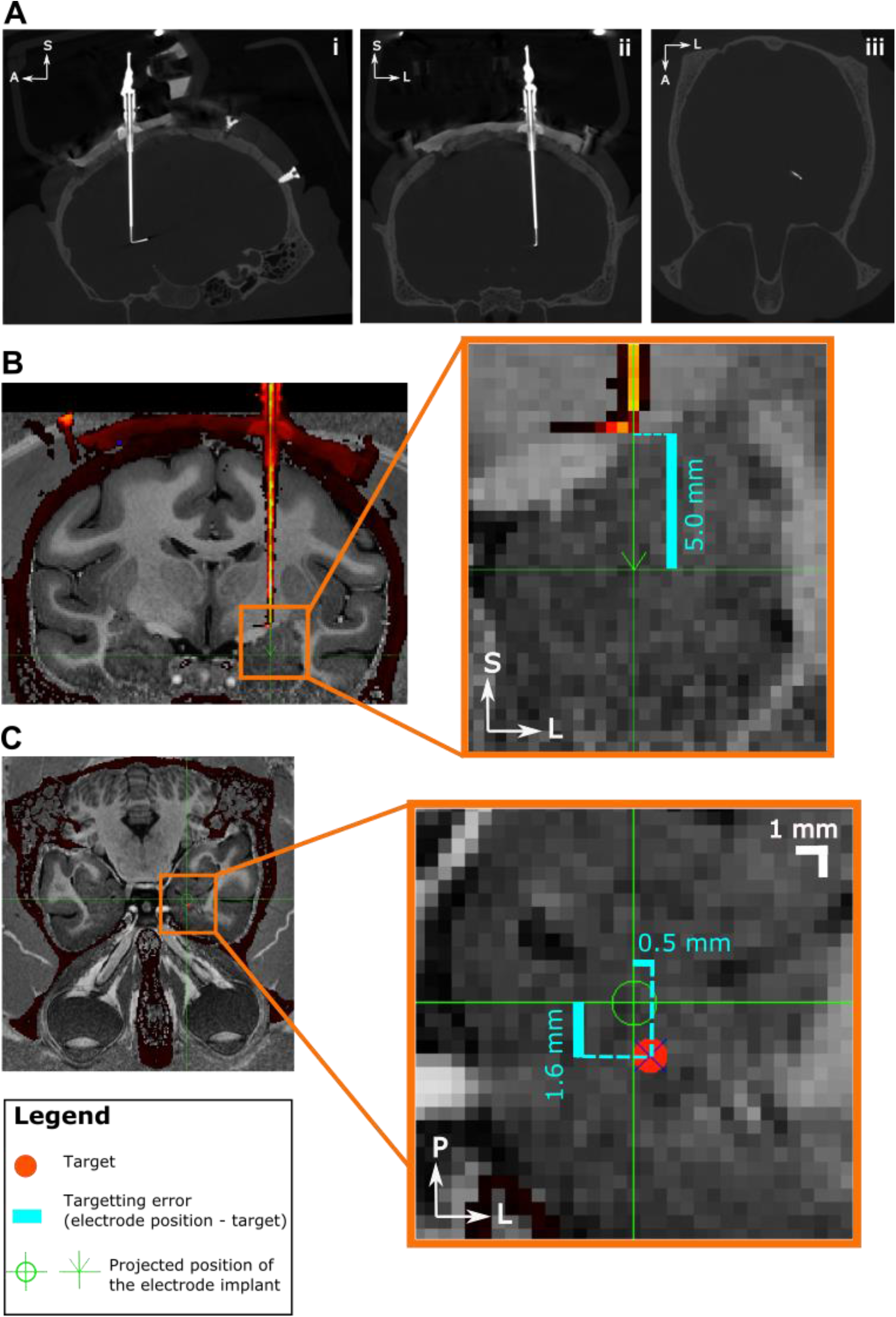
Targeting error for second implantation for subject Mi. **A.** Rotated Sagittal (i) view, to achieve an in-line image of the electrode in the post-implantation CT scan showing the bending of the microwire tips. Coronal (ii) view of the electrode, and the in-line transverse (iii) view showing the bent microwire tips. B) Bending of the microwires can be seen in the coronal view of the intensity thresholded image of the CT, co-registered with subject MRI. This is used for assessment of targeting error in the superior-inferior axis by projecting the electrode position in this axis. Anchor bolt and electrode is shown in red in the intensity thresholded image. **C.** Transverse view of the projected position of the electrode tip to enable targeting error measurement in the two axes (anterior-posterior and medial-lateral axes). Skull is shown as dark red. Insets: Magnified segments of MRI images to locate electrode tip and quantify targeting error. Actual electrode shown intensity thresholded in red pixels. Projected electrode position is shown in green. Figure legend describes inset graphic details. S = Superior; I = Inferior; A = Anterior; P = Posterior; L = Left; R = Right.

**Supplementary Table 1:** Inventory of tools and software

| ***Product*** | ***Manufacturer*** | ***Notes*** |
| --- | --- | --- |
| Anchor bolt | Ad-Tech Medical | Product number: LSBK2-BX-04 |
| Placement wrench for LSB Bolts (Anchor bolts) | Ad-Tech Medical | Product number: LSB-PWL-2.4-6X |
| Stylet/ 20-gauge Obturator | Ad-Tech Medical | Product number: OB-20-190X |
| Drill Kit | Ad-Tech Medical | DDK2-2.4-16X |
| Duo-Link Universal | BISCO | Dental cement |
| Kwik-Sil | World Precision Instruments | Low Toxicity Silicone Adhesive |
| Titanium Bone Screws | Graymatter Research | An assortment of different lengths (3-7 mm) was used depending on the skull thickness. |
| Brainsight Vet | Rogue Research Inc. | This toolkit included the computer trolley, Vicra position sensor, surgical chair and clamp, Freeguide arm, subject and pointer tracker, and assortment of tools |
| Microwire Brush Array (MBA) | Microprobes Inc. | ▪ Metal type was Platinum-Iridium (Pt-Ir).<br>▪ 32 and 64 channels were used in different implantations.<br>▪ Microwires extended 2 to 5 mm past the microfil tube. |
|  |  | Flexible cable between the MBA shaft and connectors was used<br>External ground wire was included in all but one implantation. Otherwise a stainless steel tube replaced the microfilm tube and was used as the ground connection |
| Guide tube | MicroLumen Inc. |  |
| 3D Slicer | The Slicer Community | This medical imaging software (Ver 4.10.0) was used for image co-registration and surgical planning. |
| Offline Sorter Plexon | Plexon | Software was used to perform offline sorting of the single units presented. |
| Fusion 360 | Autodesk Inc. | This software was used in designing of the microdrive. |
| Form 2 SLA | Formlabs | This 3D printer was used to fabricate the microdrive design in house. Clear V4 Resin (FLGPCL04) material was used for fabrication. Printing details are addressed in Appendix A. |
| Harris SBSKPOP Stay-Brite Silver Bearing Solder Kit | The Harris Product Group | This liquid flux was applied at the junction of the drive screw and the nut to ensure proper soldering of the two pieces. |
| AM400 | Renishaw plc | Metal 3D printer used for cranial cap of subject Ma |
| Uncoated High-Speed Steel General Purpose Tap, Taper Chamfer, 8-40 Thread Size, 3/4" Thread Length | McMaster-Carr | Part number: 2595A525<br>This tool was used to tap the body of the Microdrive which was screwed onto the anchor bolt. |
| Uncoated High-Speed Steel General Purpose Tap, Taper Chamfer, M1.6 x 0.35 mm Thread, 5/16" Thread Length | McMaster-Carr | Part number: 26015A631<br>This tool was used to tap the holes surrounding the electrode clamp groove in the Microdrive electrode clamp piece.<br>In addition it was used to tap the Microdrive body for securing the M1.6 screw for the Microdrive cover |
| Uncoated High-Speed Steel General Purpose Tap, Taper Chamfer, M2 x 0.4 mm Thread, 7/16" Thread Length | McMaster-Carr | Part number: 8305A77<br>This tool was used to tap the main body of Microdrive electrode camp which mated the drive screw through it, allowing for lowering and raising of the electrode clamp stage. |
| T-Handle Tap Wrenches | McMaster-Carr | Part number: 2544A1<br>This tool was used for hand tapping all threads in the Microdrive. |
| Super-Corrosion-Resistant<br>316 Stainless Steel Socket<br>Head Screw, M2 x 0.40 mm<br>Thread, 18 mm Long | McMaster-Carr | Part number: 92290A850<br>This was used as the drive screw to<br>lower and elevate the electrode clamp<br>in the Microdrive. |
| Super-Corrosion-Resistant<br>316 Stainless Steel Thin Hex<br>Nut, M2 x 0.4 mm Thread,<br>1.2 mm High | McMaster-Carr | Part number: 93935A305<br>This nut was fixed at the bottom of the<br>Microdrive drive screw to allow for<br>rotation in place to lower and elevate<br>the electrode clamp. Silver bearing<br>solder kit liquid flux (see above) was<br>used to ensure appropriate soldering of<br>the stainless steel screw and nut |
| Super-Corrosion-Resistant<br>316 Stainless Steel Socket<br>Head Screw, M1.6 x 0.35<br>mm Thread, 6 mm Long | McMaster-Carr | Part number: 92290A599<br>Two of these screws were used as<br>electrode clamping screws on the<br>Microdrive.<br>One of this screw type was used to<br>secure the Microdrive cover of the body<br>of the Microdrive |
| Microdrive drawings and<br>CAD files | Borna Mahmoudian, Hitarth Dalal, and Dr.<br>Julio Martinez-Trujillo | https://doi.org/10.5281/zenodo.6877605<br>This link provides the STEP and STL<br>files needed for 3D printing or<br>modification of the microdrive design.<br>As well it includes the drawing of the<br>microdrive in PDF format. |

